# Microbial GAIN domains undergo autoproteolysis and enable release of diverse cell surface associated proteins

**DOI:** 10.64898/2026.05.12.724683

**Authors:** Anna P. Brogan, David Z. Rudner

## Abstract

Adhesion G Protein-Coupled Receptors (aGPCRs) transduce mechanical stimuli across the cytoplasmic membrane in eukaryotes. These receptors contain extracellular <u>G</u>PCR <u>A</u>utoproteolysis <u>IN</u>ducing (GAIN) domains that undergo autoproteolysis but maintain stable association of their cleavage products. A diverse set of adhesion domains appended to the GAIN domain bind surface ligands on neighboring cells or the extracellular matrix. Shear force is thought to disrupt the interaction between the cleavage products exposing a tethered agonist that triggers GPCR signaling. Here, we report that GAIN domains are broadly conserved among bacteria and archaea. The microbial domains lack strong sequence conservation to their eukaryotic counterparts, but are predicted to adopt a similar fold. We demonstrate that these <u>M</u>icrobial <u>A</u>utoproteolysis <u>IN</u>ducing (MAIN) domains undergo autoproteolysis both in vitro and in vivo, using conserved catalytic residues. Furthermore, proteolysis occurs in a conserved β-turn that allows stable non-covalent interactions between the cleavage products. MAIN domains are tethered to the cell envelope of bacteria and archaea and are fused to diverse sets of adhesion and enzymatic domains. Many of the same adhesion domains are appended to both MAIN and GAIN domains, suggesting these protein families share a common origin and function. We propose that MAIN domains allow microbes to release proteins from their cell surface in response to shear force, enabling broader nutrient scavenging, intoxication of neighboring cells, and dispersal through surface detachment.

## INTRODUCTION

G-protein coupled receptors (GPCRs) are the largest family of cell surface receptors in eukaryotes. In response to external stimuli, these receptors catalyze the activation of associated G-proteins that mediate intracellular responses (1). While most GPCRs respond to chemical stimuli, a subset, known as adhesion GPCRs (aGPCRs), respond to mechanical stimuli (2–4). Like all family members, aGPCRs contain seven transmembrane (7TM) segments that undergo a conformational change in response to stimulus to activate their associated G-protein. The defining feature of the adhesion family is the presence of an extracellular <u>G</u>PCR <u>A</u>utoproteolysis <u>IN</u>ducing (GAIN) domain (**Fig. 1A**) that is fused to the 7TM signaling domain. GAIN domains undergo constitutive autoproteolysis in a loop between two β-strands (5), but the cleavage products remain non-covalently associated through an uninterrupted β-sheet. Diverse adhesion domains that bind surface ligands on neighboring cells or the extracellular matrix (**Fig. 1A**) are appended to the GAIN domain. The human genome encodes 33 aGPCRs. Each contains a GAIN domain fused to a large ectodomain with multiple adhesion domains and intrinsically disordered regions (6). Binding of these domains to surface ligands or to the matrix enables force-sensing by the aGPCR (7). A pulling or shear force is thought to be transduced to the GAIN domain causing disassociation of its cleavage products. Removal of the ectodomain reveals a β-strand known as the Stachel sequence or tethered peptide (8, 9) that is fused to the 7TM domain. The tethered peptide functions as an agonist, inducing GPCR signaling (2). Thus, aGPCRs function as mechanosensors that signal in response to mechanical rather than chemical stimuli.

**Figure 1.**
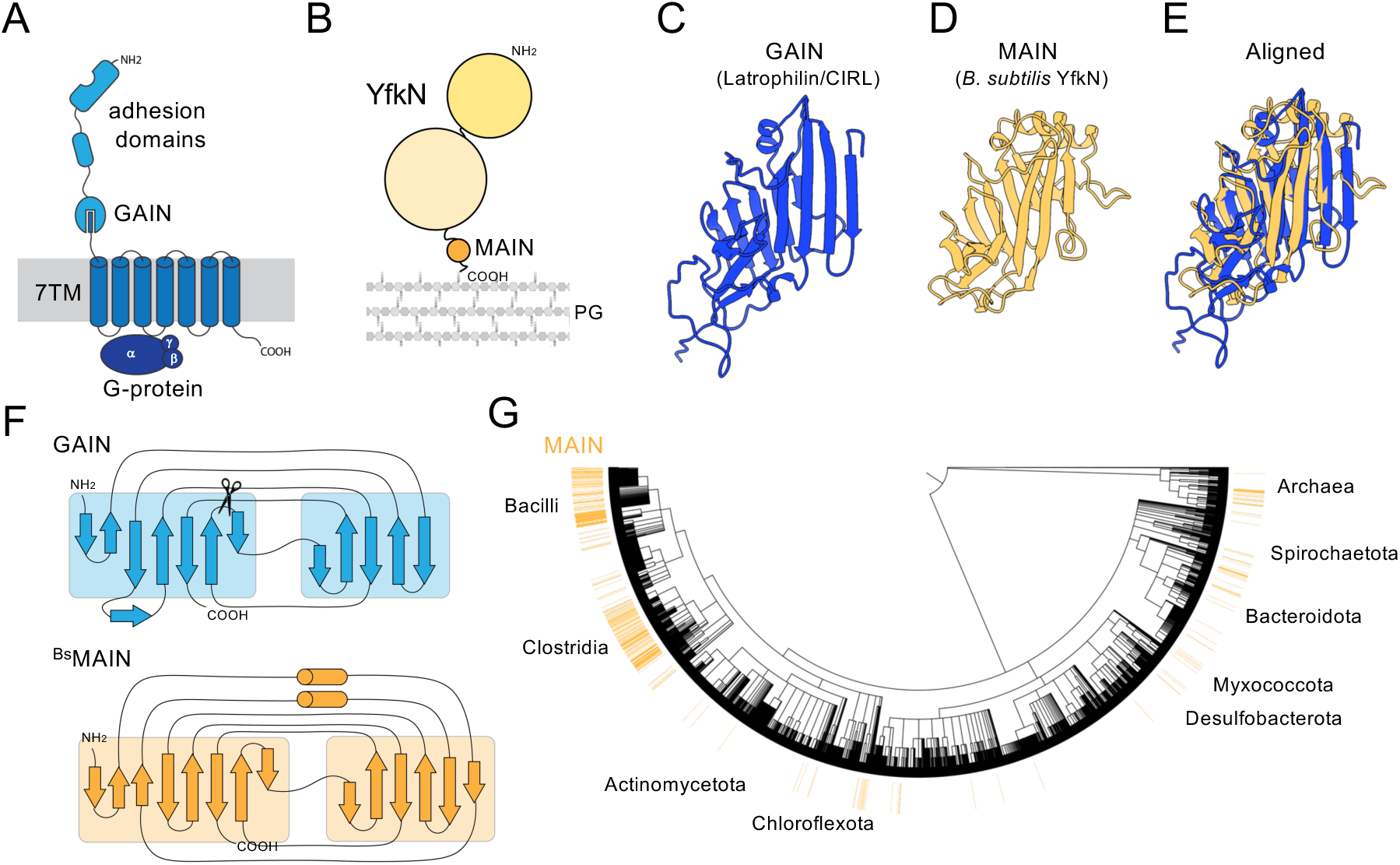
MAIN domains are widespread among bacteria and archaea. **(A)** Schematic of a prototypical adhesion GPCR. Grey represents the membrane. **(B)** Schematic of *B. subtilis* YfkN. The protein is covalently anchored to the peptidoglycan and contains a MAIN domain and two enzymatic domains. **(C)** Crystal structure of the GAIN_B_ domain from Latrophilin/CIRL (pdb:4dlq). **(D)** Alphafold2-predicted structure of the MAIN domain from *B. subtilis* YfkN. **(E)** Structural alignment by TM-align of the structures from (B) and (C). **(F)** Protein topology maps of Latrophilin/CIRL’s GAIN_B_ domain and YfkN’s MAIN domain as generated by PDMsum. Scissor depicts the site of autocleavage. **(G)** Phylogenetic tree built from 5,767 representative taxa. The amino acid sequence of ^Bs^MAIN was used to query the Refseq database and the resulting species containing a MAIN domain are highlighted in orange.

In addition to their role in aGPCR signaling, GAIN domains are found in a larger protein family of cell surface receptors, including all five human polycystic kinase disease (PKD) proteins (10). This protein family is conserved in primitive eukaryotic ancestors (5, 10), suggesting that the GAIN domain constitutes an ancient and broadly conserved mechanosensory domain.

We recently reported that bacteria encode an autoproteolytic domain called the SEAL domain that functions in a mechanotransduction pathway that is distinct from but resembles the role of the GAIN domain in aGPCR signaling (11). To uncover additional mechanosensory domains in prokaryotes, we searched for GAIN-like proteins in bacterial and archaeal genomes. We report that <u>M</u>icrobial <u>A</u>utoproteolysis <u>IN</u>ducing (MAIN) domains that are homologous to GAIN domains are encoded in many archaea and bacteria. We show that MAIN domains undergo autoproteolysis in a manner similar to GAIN domains and remain non-covalently associated after cleavage. Interestingly, MAIN domains are fused to diverse enzymatic and adhesion domains, several of which are also appended to GAIN domains. However, rather than functioning in transmembrane signaling, MAIN domains are tethered to the microbial cell surface with their enzymatic and adhesion domains facing the extracellular environment. Our findings support a model in which MAIN domains allow microbes to release hydrolytic enzymes, toxins, and adhesion domains from their cell surface in response to shear forces. We conclude that GAIN domains and their role in force-sensing are conserved throughout all domains of life and likely share a common origin.

## RESULTS

### GAIN domains are widespread throughout bacteria and archaea

The lack of reported microbial GAIN domains prompted us to investigate whether this domain or a structurally similar one exists in bacteria or archaea. Using the crystal structure of the GAIN domain from the Latrophilin/CIRL1 aGPCR (**Fig. 1C**) (5), we queried the structural homology search server Foldseek (12). Strikingly, several bacterial proteins with a highly similar fold were identified, including a strong hit in a protein (YfkN) encoded in *Bacillus subtilis* (**Fig. 1D**). The GAIN domain consists of an α-helical rich GAIN_A_ and a β-sandwich GAIN_B_ domain, which is the site of autoproteolysis. The AlphaFold2-predicted structure of the *B. subtilis* domain aligned to the Latrophilin/CIRL1 GAIN_B_ domain with a TM-align score (13) of 0.63 and a root mean squared deviation of 3.42 Å (**Fig. 1E, Fig. S1**). In addition, the topology map of the YfkN domain was nearly identical to GAIN_B_, supporting the idea that they have a common fold and likely share a common ancestor (**Fig. 1F**). Due to their similarity, we refer to these and homologous microbial domains as <u>M</u>icrobial <u>A</u>utoproteolysis <u>IN</u>ducing (MAIN) domains.

*B. subtilis* YfkN contains a signal peptide, predicted phosphodiesterase and nucleotidase domains, followed by a MAIN domain, and a C-terminal sortase recognition sequence. This sequence functions to covalently anchor the secreted protein to the cell wall via the activity of the enzyme sortase (**Fig. 1B**) (14). YfkN’s enzymatic domains have been shown to cleave cyclic nucleotides in vitro (15, 16), but the physiological function of the protein is unknown. Our bioinformatic analysis indicates that MAIN domains are found throughout bacteria and archaea with various protein architectures. They are predominantly enriched in Bacillota including Bacilli and Clostridia, but also found in Bacteroides, Spirochetes, and Archaea among other phyla (**Fig. 1G**). The cell surface anchoring of YfkN is characteristic of MAIN domain-containing proteins throughout bacteria and is discussed below.

### MAIN domains are autoproteolytic

In addition to the predicted structural homology and fusion to large ectodomains, MAIN and GAIN domains share some sequence similarity. The autocatalytic site and adjacent sequences are particularly well conserved (**Fig 2A, 2B, Fig. S3, S4**). GAIN domains undergo autoproteolysis using a conserved HX(S/T) motif, where X represents a hydrophobic amino acid. Briefly, the histidine deprotonates the serine or threonine residue, enabling attack on the adjacent amide bond to form an unstable tetrahedral intermediate. The protein then undergoes an N-O acyl shift, forming an ester intermediate, which is readily hydrolyzed by water to create two protein fragments (17, 18). It has been shown that mutating the catalytic residue to alanine abolishes autoproteolysis, and that it can be restored by mutating the serine or threonine interchangeably (18). The HXS motif is among the most highly conserved regions of bacterial MAIN domains (**Fig. 2A, Fig. S4**). To investigate whether *B. subtilis*’ MAIN domain undergoes autoproteolysis, we fused a hexa-histidine tag to the N-terminus of the MAIN domain and a SUMO protease tag to the C-terminus (His-^Bs^MAIN-SUMO). Upon expression in *Escherichia coli* and purification on Ni^2+^-agarose, we observed three polypeptides. The largest corresponded to the size of the full-length fusion protein. The sizes of the smaller two polypeptides were consistent with autocleavage products (**Fig. 2C**). Importantly, similar amounts of the two cleavage products were co-purified despite the fact that only one of them had a His-tag, suggesting that they form a stable complex after autocleavage. Autoproteolysis was abrogated by mutation of serine-1346 to alanine (S1346A) and restored by mutation to threonine (S1356T) (**Fig. 2C**). Size exclusion chromatography of the purified His-^Bs^MAIN-SUMO and the S1346A variant revealed that the autocleavage products and full-length protein have identical retention times (**Fig. 2D, 2E**). Collectively, these results argue that the mechanism of autoproteolysis is conserved between MAIN and GAIN domains, and indicate that their cleavage products remain non-covalently associated.

**Figure 2.**
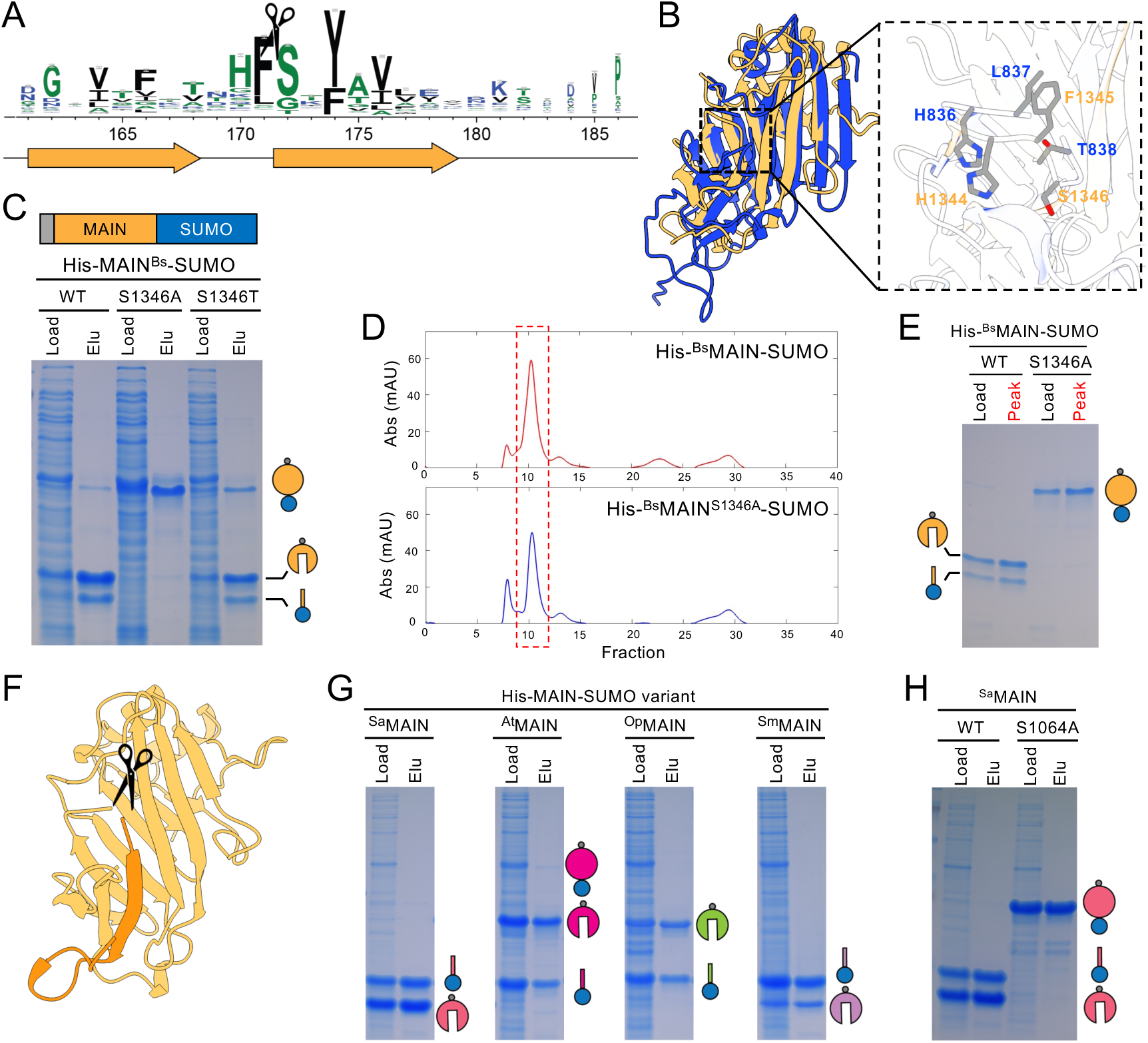
MAIN domains are autoproteolytic and remain noncovalently associated after cleavage. **(A)** Sequence logo determined from MAIN homologs in Bacillota. The autocleavage site is highlighted by a scissor. The approximate location of beta strands (arrows) and loops (lines) are depicted below. **(B)** Structural alignment (Tmalign) of Latrophilin’s GAIN domain (pdb: 4dlq) and the Alphafold2 prediction of YfkN’s GAIN domain. The inset shows the putative catalytic residues in the MAIN domain (orange) aligned with the conserved catalytic HX(S/T) residues in the GAIN domain (blue) **(C)** (top) Schematic of the His-MAIN-SUMO construct used for expression and purification. (bottom) Representative Coomassie-stained SDS-polyacrylamide gel of His-MAIN-SUMO wild-type and the indicated variants expressed and purified from *E. co*li. The load and elution (Elu) from the Ni^2+^-NTA resin are shown. Schematics depict full length and cleavage products. **(D)** Size exclusion chromatograms depicting the A280 traces for His-^Bs^MAIN-SUMO and His-^Bs^MAIN^S1346A^-SUMO. **(E)** Coomassie-stained gel of the input (Load) and peak fractions (Peak) (boxed in red) from (D). **(F)** Alphafold2-predicted structure of the autocleaved fragments of ^Bs^MAIN. The C-terminal fragment (dark orange), the N-terminal fragment (light orange), and the cleavage site (scissors) are shown. **(G)** Coomassie-stained gels of His-MAIN-SUMO variants from *Streptococcus agalactiae* (Sa), *Acetovibrio thermocellus* (At), *Oceanobacillus polygoni* (Op), *Streptococcus mitis* (Sm), and *Pseudobacteroides cellulosolvens* (Pc). Load and elution are shown. **(H)** Coomassie-stained gel of His-^Sa^MAIN-SUMO wild-type and catalytic mutant.

To determine the autoproteolytic cleavage site, we used Edman degradation of the purified His-^Bs^MAIN-SUMO protein. Cleavage occurred between the phenylalanine and serine residues of the HFS motif, at the same position where GAIN domains undergo autoproteolysis (18) (**Fig. 2F, Fig. S5**). Cleavage in this loop of the β-sheet retains extensive hydrogen bonding in the undisrupted β-sandwich, explaining how the cleavage products remain associated. Nearly all MAIN domains tested, including those from *Streptococcus agalatiae*, *Acetovibrio thermocellus*, *Oceanobacillus polygoni*, and *Streptococcus mitis* purified from *E. coli* as associated cleavage products, suggesting that they are also autoproteolytic (**Fig. 2G**). Substitution of the putative catalytic serine to alanine in the *S. agalactiae* MAIN domain abolished autocleavage (**Fig. 2H**). We conclude that bacterial MAIN domains undergo autoproteolysis identically to GAIN domains.

### MAIN domains are cell surface anchored, surface exposed, and capable of release

To investigate whether YfkN’s MAIN domain undergoes autoproteolysis in vivo, we inserted a hexa-histidine tag in the unstructured region that connects its two enzymatic domains (**Fig. 3A**) and placed the sandwich-fusion under the control of a strong IPTG-regulated promoter. To ensure that there were sufficient levels of sortase to attach His-YfkN to the cell wall, we fused the *yhcS* gene, encoding sortase, to a xylose-regulated promoter. Both expression constructs were inserted at non-essential loci in the *B. subtilis* genome. *B. subtilis* cells expressing both proteins were collected during growth in exponential and stationary phases and lysates were prepared using lysozyme followed by the addition of SDS-sample buffer. Immunoblot analysis revealed that His-YfkN was more abundant in stationary phase than exponential growth, likely because the protein was shed along with the peptidoglycan during growth (**Fig. S6**). Accordingly, all experiments were performed 4 hours after entry into stationary phase. The immunoblot in Figure 3B shows His-YfkN and the catalytic mutant His-YfkN^S1346A^. In support of the model that the MAIN domain undergoes autoproteolysis in vivo, the catalytic mutant migrated more slowly than the wild-type protein (**Fig. 3B**). A similar size shift was observed using ALFA-tagged YfkN (**Fig. S7**). It is noteworthy that His-YfkN^S1346A^ migrated as a doublet, while His-YfkN was clearly a single band (**Fig. 3B**). We wondered if the doublet corresponded to membrane-anchored and PG-anchored His-YfkN^S1346A^. Prior to attachment to the cell wall, sortase substrates are anchored in the membrane by a C-terminal transmembrane segment (19). Sortase cleaves off this TM segment and attaches the client protein to the stem peptide of the peptidoglycan precursor lipid II. Polymerization of this modified precursor incorporates the substrate into the cell wall (**Fig. S8**) (20). Thus, sortase substrates often migrate as doublets: membrane-anchored and wall-anchored (21). If His-YfkN undergoes constitutive autoproteolysis in vivo, we would not expect to see a doublet of the wild-type protein, since the C-terminal region where the modification occurs would not be covalently associated with the His-tagged YfkN cleavage product. To investigate this hypothesis, we treated cells with lysozyme in hypertonic medium and collected the soluble cell wall fraction and the protoplast pellet. The smaller His-YfkN^S1346A^ band was present in the cell wall fraction, while the larger band was present in the membrane fraction (**Fig. 3C**). By contrast, the wild-type His-YfkN band was present in both the wall and membrane fractions, as hypothesized (**Fig. 3C**). These results indicate that YfkN undergoes autoproteolysis in vivo prior to sortase-mediated attachment to the cell wall and that the cleavage products remain non-covalently associated and anchored to the peptidoglycan meshwork.

**Figure 3.**
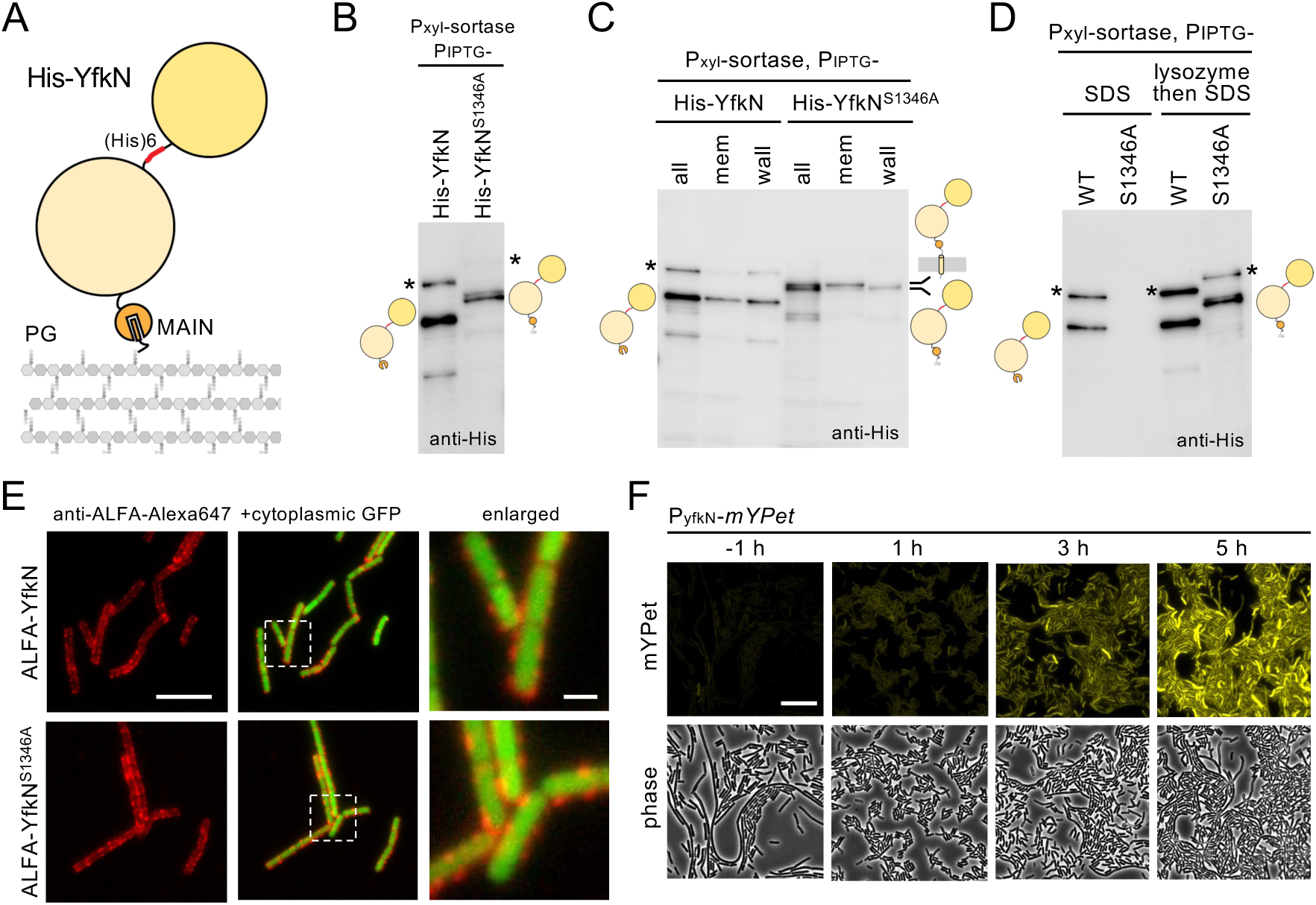
YfkN is anchored to the cell surface and undergoes autoproteolysis *in vivo*. **(A)** Schematic of His-YfkN. YfkN is anchored to the peptidoglycan (PG) at its C-terminus by sortase, followed by its MAIN domain (orange), 5’ nucleotidase (light yellow), and phosphodiesterase (dark yellow) domains. The (His)_6_ tagged was inserted between the two enzymatic domains. **(B)** Immunoblot of stationary phase *B. subtilis* cells constitutively expressing His-YfkN or the catalytic mutant S1346A. Schematics of His-YfkN variants are depicted adjacent to relevant bands on the blot. Anomalous migrating species are indicated (*). **(C)** Immunoblot of His-YfkN fractionation. Cells were protoplasted and membrane and wall fractions were analzyed. **(D)** Immunoblot of SDS-released His-YfkN variants. **(E)** Representative fluorescence micrographs of the indicated *B. subtilis* stationary phase cells expressing cytoplasmic GFP probed with anti-ALFA-Alexa647. The boxed regions in the merged image were enlarged to more clearly visualize the surface-localized ALFA-YfkN. Far left scale bar indicates 5 µm and far right scale bar indicates 1.3 µm. *B. subtilis* YfkN was expressed with 500 µM IPTG and sortase was expressed with 20 mM xylose in all backgrounds (B-E) **(F)** Fluorescence and phase-contrast micrographs of cells harboring a P*yfkN*-*mYPet* reporter. Time in hours (h) before and after entrance into stationary phase are indicated. Scale bar indicates 10 µm. The grow curve associated with this experiment is shown in Figure S13.

We hypothesize that the enzymatic domains of YfkN are surface-exposed, but can be released in response to specific stimuli. To investigate whether YfkN can be released into the extracellular environment, we treated *B. subtilis* cells expressing His-YfkN or His-YfkN^S1346A^ with SDS and monitored the released proteins by immunoblot. As anticipated, the autoproteolyzed wild-type His-YfkN was released by SDS, while the membrane- and PG-anchored His-YfkN^S1346A^ proteins were not (**Fig. 3D**). In addition, we found that exposing cells to 95 °C for 5 minutes similarly resulted in release of His-YfkN’s ectodomain, while cells expressing the autoproteolysis deficient mutant did not (**Fig. S9**). Together, these results provide further support for MAIN domain autoproteolysis in vivo and indicate that YfkN’s ectodomain can be released into the environment, in these cases by non-physiological stimuli. To test whether YfkN is surface-exposed, we inserted an ALFA tag between the signal sequence and the first enzymatic domain of YfkN and expressed the fusion protein under IPTG control. We monitored surface-associated ALFA-YfkN in late stationary phase using a fluorophore-conjugated anti-ALFA nanobody that cannot cross the cell wall (**Fig. S10**). As anticipated, cells expressing ALFA-YfkN and ALFA-YfkN^S1346A^ were readily labeled on their cell surface (**Fig. 3E**).

Peptidoglycan is turned over as cells grow and divide. Accordingly, YfkN would be constantly released into the medium with the PG cleavage products if expressed throughout exponential growth, obviating the need for the MAIN domain. To determine when *yfkN* is transcribed, we fused the *yfkN* promoter to *mYPet* and inserted this transcriptional reporter at a non-essential locus in the *B. subtilis* genome. We assayed *yfkN* promoter activity during growth in LB and low phosphate defined medium (LPDM), a medium that has previously been shown to increase production of YfkN (15, 22). mYPet fluorescence increased in late stationary phase, consistent with the idea that YfkN is only expressed and anchored to the cell wall at a time when the cells have largely stopped growing (**Fig. 3G**). As reported previously, we observed stronger induction in LPDM than LB (**Fig. S11**) (15). In sum, our data indicate that YfkN is produced when cells stop growing, the secreted protein undergoes autoproteolysis but the cleavage products remain associated, and the protein is then attached to the cell wall and becomes surface exposed. We propose that the enzymatic domains can act on substrates adjacent to the surface and, if released by shear force, can access substrates further away from the cell.

### MAIN domains are found in diverse protein architectures

To survey the breadth of domains associated with a MAIN domain, we compiled 15,000 MAIN-containing proteins across bacteria and archaea and annotated known Pfam domains within their protein sequences using PfamScan (23) (**Table S1, S2)**. In a complementary approach, we clustered the 15,000 homologs and used AlphaFold2 to predict the structure of 2,843 unique MAIN domain-containing proteins. We subsequently used Domain Parser for Alphafold Models (DPAM) (24) to annotate known domains with structural similarity (**Fig. S12**). Highlighting the incredible diversity of proteins which contain a MAIN domain, we identified 350 distinct Pfam domains from the sequences and 3,874 distinct ECOD (Evolutionary Classification of protein Domains) domains from the predicted structures (**Table S3, S4**). Several observations emerged from this analysis. First, virtually all the proteins contain a signal peptide as predicted by SignalP (25), indicating that the proteins are secreted across the cytoplasmic membrane. Second, the vast majority of MAIN-domain containing proteins contain a sortase recognition sequence, cell wall anchoring domains, or an S-layer homology domain, which binds glycopolymers that are covalently attached to the cell wall (26). The presence of these domains indicates that MAIN domain-containing proteins are typically anchored to the cell surface. Third, most MAIN domain-containing proteins are large by bacterial standards, the average protein molecular weight among the 15,000 homologs was 158 kDa. Finally, the other domains associated with MAIN domains could be grouped into two categories: those that have predicted roles in adhesion, and those that have enzymatic function. MAIN-domain containing proteins often possess both types of domains, and often with multiple copies of each (**Fig. 4A**). Several of the adhesion domains that are associated with MAIN domains are also found on human aGPCRs, including Ig-like, Cadherin-like, Fibronectin-like, Leucine Rich Repeats, and lectins (**Fig. 4A, S13**). Interestingly, a subset of MAIN-domain containing proteins are fused to putative toxins in their ectodomains (**Fig. 4A**).

**Figure 4.**
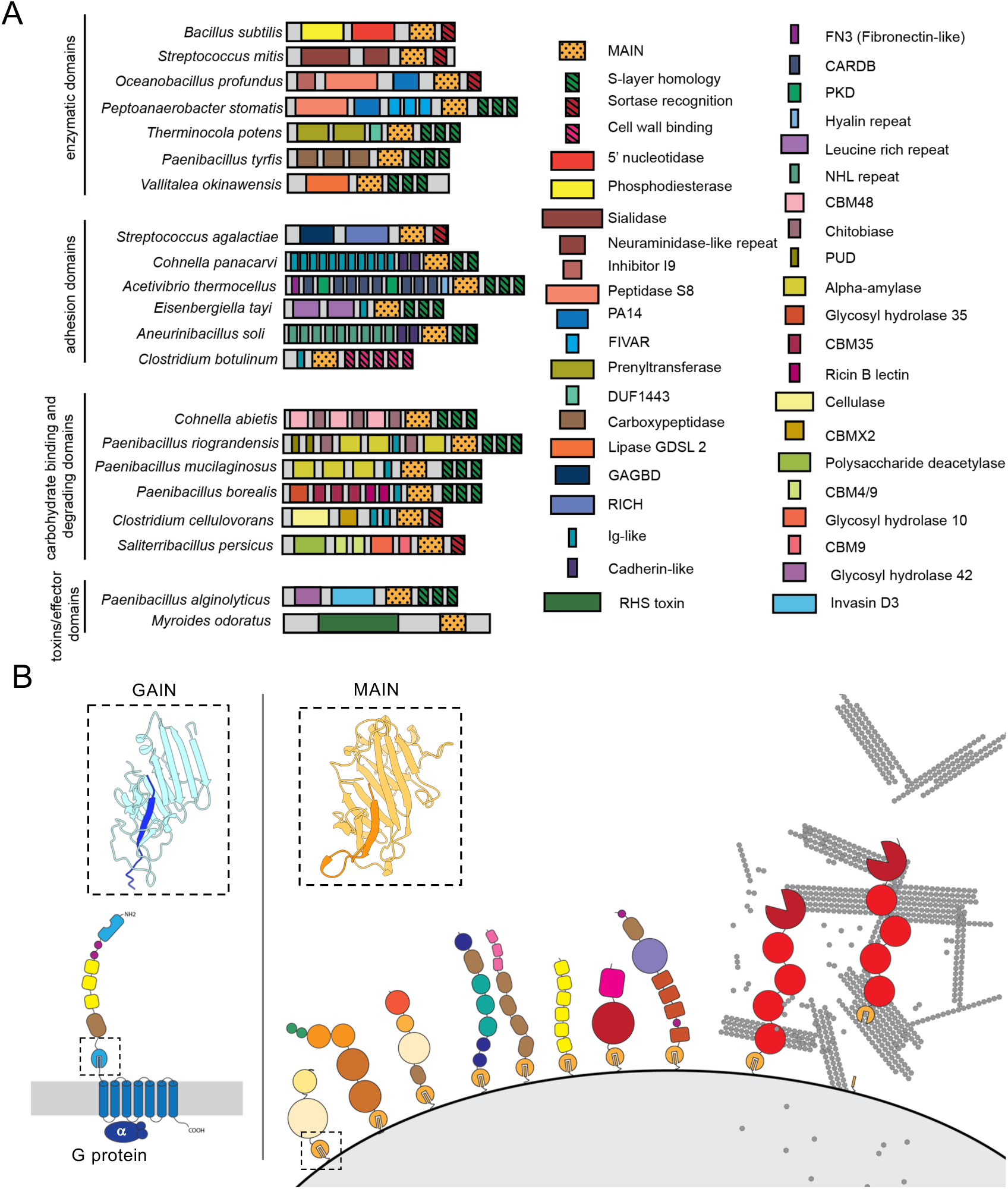
Schematic of MAIN and GAIN domain-containing proteins on the cell surface. **(A)** Schematic of representative MAIN domain containing proteins from diverse bacteria. **(B)** Schematic of a prototypical aGPCR and inset of the crystal structure of Latrophilin/CIRL’s GAINB domain with N-terminal (light blue) and C-terminal (dark blue) cleavage products. The schematic on the right depicts examples of MAIN domain containing proteins. These proteins are anchored onto microbial cell surfaces and display large and diverse enzymatic and adhesion domains. The example on the far-right depicts a cellulose-binding and degrading protein released by interaction with cellulose.

A GO enrichment analysis of the Pfam domains associated with MAIN domains revealed several enzymatic functions, the most prominent of which was hydrolase activity, particularly on glycosidic bonds (**Fig. S14**). Furthermore, many of these proteins contain both glycoside hydrolase domains and carbohydrate-binding lectins (**Fig. 4A**), a common arrangement for carbohydrate degrading enzymes (27). Notably, many of these MAIN-domain containing proteins are encoded by bacterial species in the human and soil microbiome (**Table S2, S3**). Finally, virtually all the proteins analyzed contained a similar domain arrangement (from N to C-terminus): signal peptide, adhesion and enzymatic domains, MAIN domain, and cell surface anchoring domain. These findings support the hypothesis that MAIN domains are autoproteolytic and function to release cell surface associated enzymatic, adhesion, and toxin domains in response to shear to enable broader nutrient scavenging, dispersal, or intoxication of neighboring cells (**Fig. 4B**).

### Diverse MAIN domains are cell surface-anchored and autoproteolytic

To extend our analysis to the diverse set of proteins containing autoproteolytic MAIN domains, we engineered cells to express and anchor heterologous substrates onto the *B. subtilis* cell wall. To do so, we introduced the N-terminal YfkN signal sequence and C-terminal cell wall anchoring sequence in place of those present on candidate proteins (**Fig. S15A**). In addition, an ALFA tag was inserted between the signal sequence and the candidate proteins. These chimeras were expressed under the control of a strong IPTG-regulated promoter alongside the xylose-regulated sortase, YhcS. We analyzed two candidate proteins: a MAIN-containing protein from *Pseudoruminococcus massiliensis* (WP_106763535) that contains a Leucine Rich Repeat and putative starch binding domain CBM26, and one from *Gracilibacillus dipsosauri* (WP_109985763) containing a cadherin-like domain, a glycosyl hydrolase 26 domain with predicted activity on xylans and mannans, and a putative mannan binding CBM35 domain. Both MAIN domains were autoproteolytic, as assayed by purification of His-MAIN-SUMO wild-type and autoproteolysis deficient mutants from *E. coli* (**Fig. S15B**). In addition, immunoblot analysis of *B. subtilis* lysates revealed size shifts consistent with in vivo autoproteolysis for both chimeras (**Fig. S15C**). Finally, immunolabeling of *B. subtilis* cells expressing the sortase anchored chimeras indicated that they are cell surface exposed (**Fig. S15D**). Of note, unlike native YfkN, both chimeras had to be expressed in a strain lacking extracellular proteases for efficient expression and cell surface exposure, likely due to non-specific proteolysis of the non-native proteins. Altogether, we conclude that diverse MAIN-domain containing proteins are autoproteolytic in vivo.

## DISCUSSION

Altogether, our findings define a new bacterial autoproteolytic domain, expand the phylogenetic diversity of GAIN-like domains, and provide a model for a new form of mechanosensing in bacteria. We show that diverse microbial proteins with MAIN domains are autoproteolytic, anchored to the cell surface, and capable of release. GAIN domains function in aGPCR signaling as mechanotransducive domains that are pulled apart in response to mechanical stimuli, allowing ectodomain release and GPCR signaling. By analogy, we propose that MAIN domain-containing proteins sense mechanical stimuli when bound to complex polysaccharides, neighboring cells, or surfaces and release their ectodomains in response to shear forces (**Fig. 4B**).

Many gut and soil bacteria encode MAIN domains that tether large ectodomains consisting of repeated adhesion domains, lectins, and glycosyl hydrolases with predicted specificity for complex polysaccharides. We hypothesize that the lectin domains bind polysaccharides derived from plant cell walls in the gut lumen or the soil and shear force results in the dissociation of the MAIN domain cleavage products. Release of the enzymatic domains enables degradation of polysaccharides and nutrient scavenging at a distance (**Fig. 4B**). The force transduced upon polysaccharide binding is reminiscent of sigma/anti-sigma signaling by SigI/RsgI systems in Clostridial species. These RsgI anti-sigma factors contain extracytoplasmic SEA-like domains called SEAL domains that undergo autoproteolysis but the cleavage products remain stably associated. A carbohydrate-binding domain at the extreme C-terminus of RsgI binds cellulose or other polysaccharides and a resulting shear force is hypothesized to pull the SEAL domain apart (11, 28). This likely triggers intramembrane proteolysis, sigma factor activation, and expression of the proteins that make up a multi-protein complex responsible for cellulose degradation, called the cellulosome (29, 30).

A subset of MAIN domains are fused to toxin domains, or domains at the bacteria-host interface. For example, the wall-anchored protein IgA FC receptor in Group B Streptococcus (*S. agalactiae*) contains a MAIN domain. The IgA FC receptor binds the Fc region of human IgA enabling this bacterium to evade immune detection (31, 32). Similar to YfkN, the MAIN domain is present at the extreme C-terminus of the IgA FC receptor, adjacent to the sortase anchoring domain. Our findings raise the possibility that the MAIN domain enables release of the receptor from the *S. agalactiae* surface when it is bound by IgA. Some MAIN domains are also fused to RHS repeat-associated toxin domains. Interestingly, RHS toxins are often found on contact-dependent inhibition (CDI) systems (33). In Gram-negative bacteria, some of these CDI toxins are anchored in the outer membrane by β-barrels and toxin release is mediated by physical contact (34). Our findings strongly suggest that the MAIN domain could mediate toxin delivery in response to cell-cell contact in a manner analogous to these Gram-negative CDI systems.

This proposed mode of toxin delivery shares striking similarities to Type V secretion via autotransporters in Gram-negative bacteria (35). Autotransporters assemble a C-terminal β-barrel translocator domain in the outer membrane. It is thought that after folding, the N-terminal domain, known as the passenger or ⍺ domain, passes through the translocator and is released by proteolytic cleavage. This cleavage can be autoproteolytic or catalyzed by a membrane-anchored protease (36–38). In some cases, after cleavage, the passenger and β-domains remain non-covalently associated (39). Moreover, the passenger domains in these systems can be highly variable and have roles in adhesion and virulence (40, 41). These similarities suggest that MAIN domains could be the Gram-positive counterpart to Gram-negative autotransporters.

Many of the same adhesion domains that are present on aGPCRs are also fused to MAIN domains in bacteria. In addition, aGPCR and MAIN-containing proteins have the same protein architecture: large N-terminal ectodomains, MAIN or GAIN domain, and cell surface anchoring domains. A key difference between the two is that release of the ectodomain is the regulated step of aGPCR signaling and alters intracellular physiology, whereas no signaling mechanisms have been associated with the C-terminal fragment of the MAIN domain that remains anchored to the cell wall. Whether these surface-anchored “tethered peptides” impact cellular physiology remains to be determined. However, it is noteworthy that aGPCRs are thought to have two modes of signaling: *cis* and *trans.* In the *cis* mode, the tethered agonist binds the 7TM domain to which it is fused, resulting in G-protein activation. In the *trans* mode, signaling occurs via interaction of the released ectodomain with an extracellular partner (4). This latter mode of signaling is analogous to the proposed release of the cleaved MAIN and associated domains from the microbial cell surface in response to force. We speculate that this represents the ancestral aGPCR signaling mode.

The finding of a conserved protein fold across all domains of life suggests an ancient function for the MAIN and GAIN domain. Both domains, as well as the mechanosensitive SEA and SEAL domains, undergo autoproteolysis in a loop between two β-strands within a β-sheet. This fold allows non-covalent association of the cleavage products via an extensive hydrogen bonding network. In response to mechanical stimuli, the cleavage products dissociate, allowing signaling in *cis* or *trans* (2, 11, 28, 42, 43). Like domains fused to MAIN and GAIN domains, SEAL and SEA domains are frequently linked to adhesion domains, often of the same fold across domains, suggesting the possibility of a similar signaling mechanism (**Fig. S16, S17**) (11). The convergent evolution of these force-sensing domains reveals a simple and robust solution to sensing and responding to mechanical stimuli.

## MATERIALS AND METHODS

### General methods

All *B. subtilis* strains were derived from the prototrophic strain PY79 (44). Unless otherwise indicated, cells were grown in LB at 37 °C. Insertion-deletion mutations were generated by isothermal assembly (45) of PCR products followed by direct transformation into *B. subtilis*. The chimeric MAIN domain-containing proteins were engineered in the strain WB600N, lacking seven extracellular proteases (46). Tables of strains, plasmids and oligonucleotide primers and a description of strain and plasmid construction can be found online as supplementary data (Supplemental Table S5, S6, S7, S8, and Text S1).

### Immunoblot analysis

For Immunoblot analysis, cells were subcultured in 3 mL LB for 2.5 hours, back-diluted into 25 mL LB containing 500 µM IPTG and 20 mM xylose at OD600 = 0.05 and grown for 6 hours (4 hours into stationary phase). 1 mL of cells normalized to OD600 = 1 was pelleted and resuspended in lysis buffer (20 mM Tris pH 7.0, 10 mM MgCl_2_ and 1mM EDTA, 1 mg/mL lysozyme, 10 µg/mL DNase I, 100 µg/mL RNase A, 1 mM PMSF) to a final OD_600_ of 20 for equivalent loading. The cells were incubated at 37 °C for 15 min followed by addition of an equal volume of Laemelli sample buffer (0.25 M Tris pH 6.8, 4% SDS, 20% glycerol, 10 mM EDTA) containing 10% β-mercaptoethanol. Samples were heated for 5 min at 95 °C prior to loading. Proteins were separated by SDS-PAGE on 7.5 % polyacrylamide gels, electroblotted onto Immobilon-P membranes (Millipore) and blocked in 5% nonfat milk in phosphate-buffered saline (PBS) with 0.5% Tween-20. The blocked membranes were probed with anti-SigA (1:10,000) (ref), anti-His (1:4,000) (GenScript), anti-ALFA (1:1,000) (Nanotag) antibodies diluted into 3% BSA in PBS with 0.05% Tween-20. Primary antibodies were detected using horseradish peroxidase-conjugated goat anti-rabbit or anti-mouse IgG (BioRad) and the Super Signal chemiluminescence reagent as described by the manufacturer (Pierce). Signal was detected using a Bio-Techne FluorChem R System.

When cell wall and membrane fractions were separated, cell pellets were washed 2 times in 1XSMM (0.5 M sucrose, 20 mM MgCl_2_, 20 mM maleic acid pH 6.5), resuspended in 50 µL SMM + 0.5 mg/mL lysozyme + 5 U Mutanolysin and incubated at room temperature for 30 minutes. The cells were monitored by light microscopy until >95% had formed protoplasts. The protoplasts were pelleted by centrifugation at 3k rpm for 10 minutes. The supernatant was collected and mixed with equal volume 2X Laemelli sample buffer containing 10% β-mercaptoethanol. The protoplast pellet was resuspended in 100 µL 1X Laemelli sample buffer containing 5% β-mercaptoethanol. Samples were further processed as described above.

For heat treatment, cell pellets were resuspended in 100 µL PBS + 1 mM PMSF and heated at 95°C for 5 minutes. The cells were pelleted (4000g/4min), the supernatant combined with 2X Laemelli sample buffer containing 10% β-mercaptoethanol, and the cell pellet was lysed as described above.

For SDS treatment with and without lysozyme, 20 mL of culture was resuspended in 1 mL PBS + 1 mM PMSF + 10 µg/mL DNase I + 100 µg/mL RNase with or without 0.5 mg/mL lysozyme. Cells were lysed via sonication and 100 µL of the sample combined with 100 µL 2X Laemelli sample buffer containing 10% β-mercaptoethanol.

### Purification of His-MAIN-SUMO variants

Expression plasmids containing His-MAIN-SUMO variants were transformed into *E.coli* BL21(DE3) Δ*tonA* cells. Transformants were sub-cultured in terrific broth (TB) + 100 µg/mL ampicillin for 3 hours then back-diluted into 25 mL TB + 100 µg/mL ampicillin to an OD600 of 0.01 and grown at 37 °C to an OD600 of 0.4. Cultures were induced with 500 µM IPTG for 3 hours at 37 °C and then harvested by centrifugation at 4000 rpm for 15 minutes. Cells were concentrated 25-fold by resuspending the pellet in 1 mL Lysis Buffer (50 mM HEPES-NaOH, 300 mM NaCl, 25 mM Imidazole, 10% (v/v) glycerol) and frozen at -80 °C. Cell pellets were thawed on ice and lysed by sonication. 25 U Benzonase (Sigma) and 1X complete protease inhibitor (Roche) were added to the cell lysate and incubated on ice for 15 min. Lysates were clarified by ultracentrifugation at 40,000 rpm for 45 minutes at 4 °C. The supernatant was added to 25 µL of Ni^2+^-NTA resin, washed with 120 bed volumes of wash buffer (20 mM HEPES-NaOH, 300 mM NaCl, 25 mM Imidazole, 10% (v/v) glycerol), and eluted in 100 µL of elution buffer (20 mM HEPES-NaOH, 300 mM NaCl, 300 mM Imidazole, 10% (v/v) glycerol). Load and Elution fractions were resolved by 12.5% SDS-PAGE gels and stained with Instant Blue (Abcam).

### Size Exclusion Chromatography

His-MAIN-SUMO and His-MAIN^S1346A^-SUMO were analyzed by size-exclusion chromatography (SEC) on a Superdex S75 column (GE Healthcare) in buffer containing 20 mM of HEPES-NaOH, pH 7.5 and 300 mM of NaCl. The load and peak absorbance (A280) fractions were diluted into equal volume 2X sample buffer containing 10% β-mercaptoethanol and resolved by SDS-PAGE using a 17.5% polyacrylamide gel and stained with Instant Blue (Abcam). Absorbance profiles were plotted using MATLAB.

### Edman Degradation

Purified His-MAIN-SUMO as described above was separated by SDS-PAGE using a 17.5% polyacrylamide gel and transferred to a PVDF membrane at 90 V for 60 min. The membrane was then washed 5X with ddH_2_O, stained with 0.02% Coomassie Brilliant blue in 40% methanol, 5% acetic acid for 30 sec, destained in 40% methanol, 5% acetic acid for 1 min, washed 3X with ddH_2_O and allowed to air dry. The bands of interest were cut from the membrane and sent for 5 cycles of Edman Degradation analysis with an ABI 494 Protein Sequencer (Tufts University Core Facility).

### Bioinformatics Analysis

A local PSI-BLAST run was separately performed on the amino acid sequences of *B. subtilis* YfkN’s MAIN domain and *H. sapiens* MUC1’s SEA domain against the RefSeq Select Protein database using an e-value of 0.05 and 5 iterations. For the MAIN domain, eukaryotic proteins were removed from the results and the remaining protein sequences retrieved (47). TaxIDs of all MAIN domain homologs were used to plot onto a phylogenetic tree of 5767 unique bacterial taxa. Tree visualization and manipulation was done using iTOL (48).

#### PfamScan

The ∼15,000 proteins identified by PSI-BLAST were annotated using PfamScan against the Pfam database release 31 (23). The output domains were organized by RefSeq accession number and the resulting domain organizations built (**Table S2, Fig. S15**). Counts of each domain found associated with the MAIN domain can be found in Table S1.

#### DPAM

MAIN domain-containing proteins identified by PSI-BLAST were clustered using MMseqs2 using a minimum sequence identity of 50% and a coverage threshold of 80%. The structures of the resulting 2,843 unique proteins were predicted using colabfold version 1.5.5 (49, 50) with the following parameters: each prediction was run with five models, using three recycles, no dropout, no templates. These runs were performed using a local installation of Colabfold running on NVIDIA A6000 GPUs managed by the HMS O2 computing cluster. MSAs were generated using MMseqs2 API server (51). The rank 1 prediction (based on pLDDT) from each AlphaFold2 run was used as an input into DPAM (24) (**Fig. S12**). The resulting output domains were organized by RefSeq accession number and the domain organizations built (**Table S3, S4**).

### Multiple Sequence Alignment

Multiple sequence alignments were generated from the results of the previously described psi-blast searches using Clustal Omega (52). NCBI taxID was used to filter the MSA for organisms from the Bacillota (Firmicute) phylum.

### Alphafold2 Predictions

Protein structures were modeled using colabfold version 1.5.5 (49, 50). Each prediction was run with five models, using three recycles, no dropout, no templates. These runs were performed using a local installation of Colabfold running on NVIDIA A6000 GPUs managed by the HMS O2 computing cluster. MSAs were generated using MMseqs2 API server (51). All predictions used paired and unpaired MSAs, and structures were unrelaxed. The highest ranked structure by pLDDT was used for structural annotation.

### Structural model visualization

The crystal structure of CIRL1/Latrophilin’s GAIN domain (pdb: 4dlq) was downloaded from the PDB. ChimeraX1.3 and Pymol 2.4.0 were used to visualize the structural models and generate images. PDBsum was used to generate protein topology maps.

### Fluorescence Microscopy

Fluorescence microscopy was performed on a Nikon Ti2-E inverted microscope equipped with Plan Apo 100x/1.45 phase contrast oil objective and a Hamamatsu ORCA-Flash4.0 V3 Digital CMOS camera. Light was delivered with the Lumencor Spectra III Light Engine containing the following filters: 390/22, 440/20, 475/28, 510/25, 555/28, 575/25, 637/12, 748/12. Images were acquired using the Nikon Elements 5.2 acquisition software. Cytoplasm was labeled with constitutively expressed GFP when indicated. Images were cropped and adjusted using Fiji. All images shown side by side were acquired, analyzed, and adjusted, identically for comparison. Images are shown at a magnification of 100x and cropped to 126.88 µm x 126.88 µm unless otherwise indicated.

### P_yfkN_-*mYPet* expression

Cells were subcultured in low phosphate defined media (LPDM) for 3 hours before back-dilution into 25 mL LPDM at OD600 = 0.05 and grown for 9 hours. 1 mL of cells was sampled every 30 minutes, the OD600 recorded, pelleted, resuspended in 50 µL LPDM, spotted onto 2% LPDM agarose, and imaged as described in fluorescence microscopy. LPDM consists of: 50 mM Tris pH 7.1, 3.03 mM (NH_4_)_2_SO_4_, 6.8 mM trisodium citrate, 3.04 mM FeCl_3_, 1 mM MnCl_2_, 3.5 mM MgSO_4_, 0.01 mM ZnCl_2_, 0.5% glucose, 0.05% Casamino acids, 10 mM L-arginine, 0.065 mM KH_2_PO_4_. Similar results were obtained using LB medium.

### Anti-ALFA Immunolabeling

Cells expressing ALFA-tagged proteins were subcultured in 3 mL LB for 2.5 hours, back-diluted into 25 mL LB containing 500 µM IPTG and 20 mM xylose at OD600 = 0.05 and grown for 6 hours into stationary phase. 1 mL of cells normalized to OD600 = 1 was pelleted, washed 1X in PBS, and labeled in PBS + 1:500 anti-ALFA-Alexa647 (Nanotag) for 15 minutes at room temperature. Cells were washed 1X in PBS and resuspended in 100 µL PBS before spotting 6.5 µL onto 2% PBS agarose pads and imaging as described in fluorescence microscopy.

## Acknowledgements

We thank Shailab Shrestha and all members of the Bernhardt-Rudner supergroup past and present for helpful advice, feedback, discussions, and encouragement, the Tufts University Core Facility (TUCF) for Edman degradation analysis, and the Microscopy Resources on the North Quad (MicRoN) core at Harvard Medical School for help with microscopy. Portions of this research were conducted on the O2 High Performance Computing Cluster, which is supported by the Research Computing Group at Harvard Medical School. Support for this work comes from the National Institute of Health Grants R35GM145299, U19 AI158028 (D.Z.R.). A.P.B. was funded in part by NSF (DGE1745303) and NIH (F31AI181098).

## Data availability

Uncropped immunoblots are included in the supplementary information. Strains, plasmids, and oligonucleotides used can be found in supplementary tables.

## Contributions

A.P.B. and D.Z.R. conceived the study, A.P.B. performed the experiments, A.P.B. and D.Z.R. performed the analyses, D.Z.R. supervised the study, A.P.B. and D.Z.R. wrote the paper.

## Declaration of generative AI and AI-assisted technologies in the manuscript preparation process

During the preparation of this work the authors used ChatGPT in order to suggest new words and alternative sentence structure. After using this tool/service, the authors reviewed and edited the content as needed and take full responsibility for the content of the published article.

---

**Figure S1.**
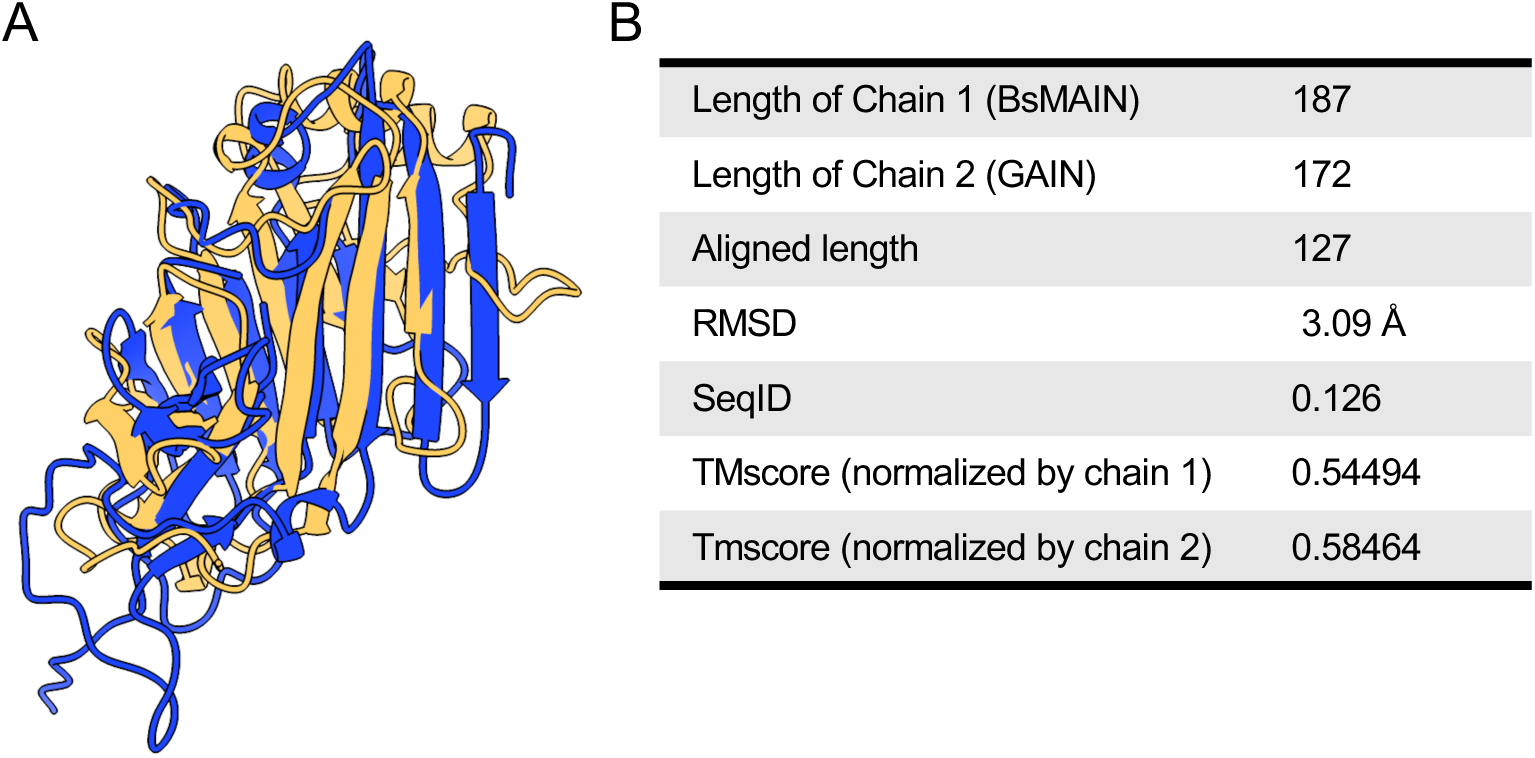
Alignment metrics for BsMAIN and Latrophilin/CIRL GAIN. **(A)** Structural alignment by TM-align of the structures from Latrophilin/CIRL1’s GAIN domain (blue) and YfkN’s GAIN domain (orange). Confidence metrics for the Alphafold2 prediction of *the B. subtilis* MAIN domain can be found in Figure S2. **(B)** Metrics from the structural alignment using TMalign.

**Figure S2.**
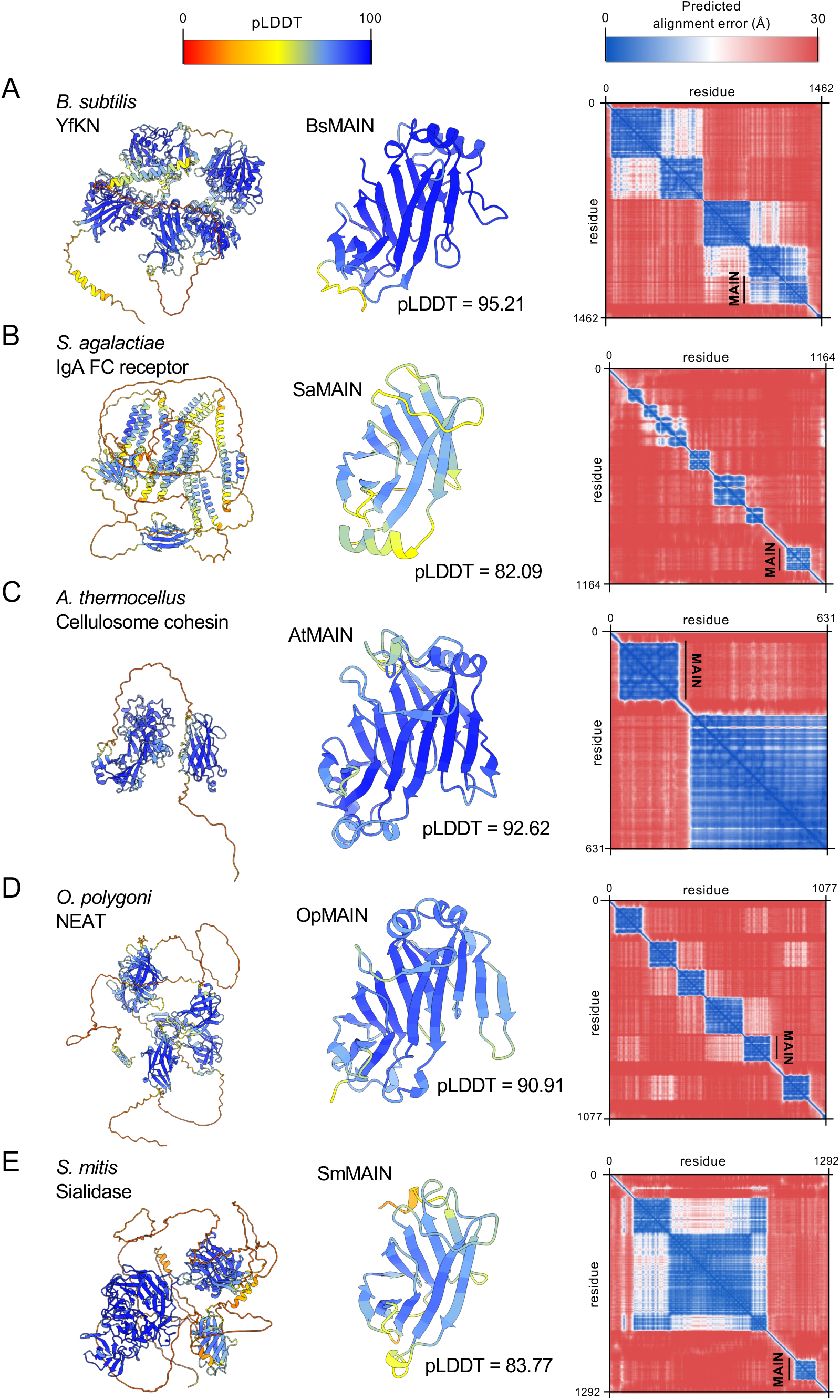
Metrics for Alphafold2 predictions. Alphafold2 predictions of full-length MAIN domain containing proteins (left) and MAIN domains (center) colored by pLDDT and predicted alignment error (pAE) plots (right) for **(A)** *B. subtilis* YfkN, **(B)** *S. agalactiae* IgA FC receptor, **(C)** *A. thermocellus* cellulosome anchoring cohesin, **(D)** *O. polygoni* NEAT transporter, and **(E)** *S. mitis* sialidase.

**Figure S3.**
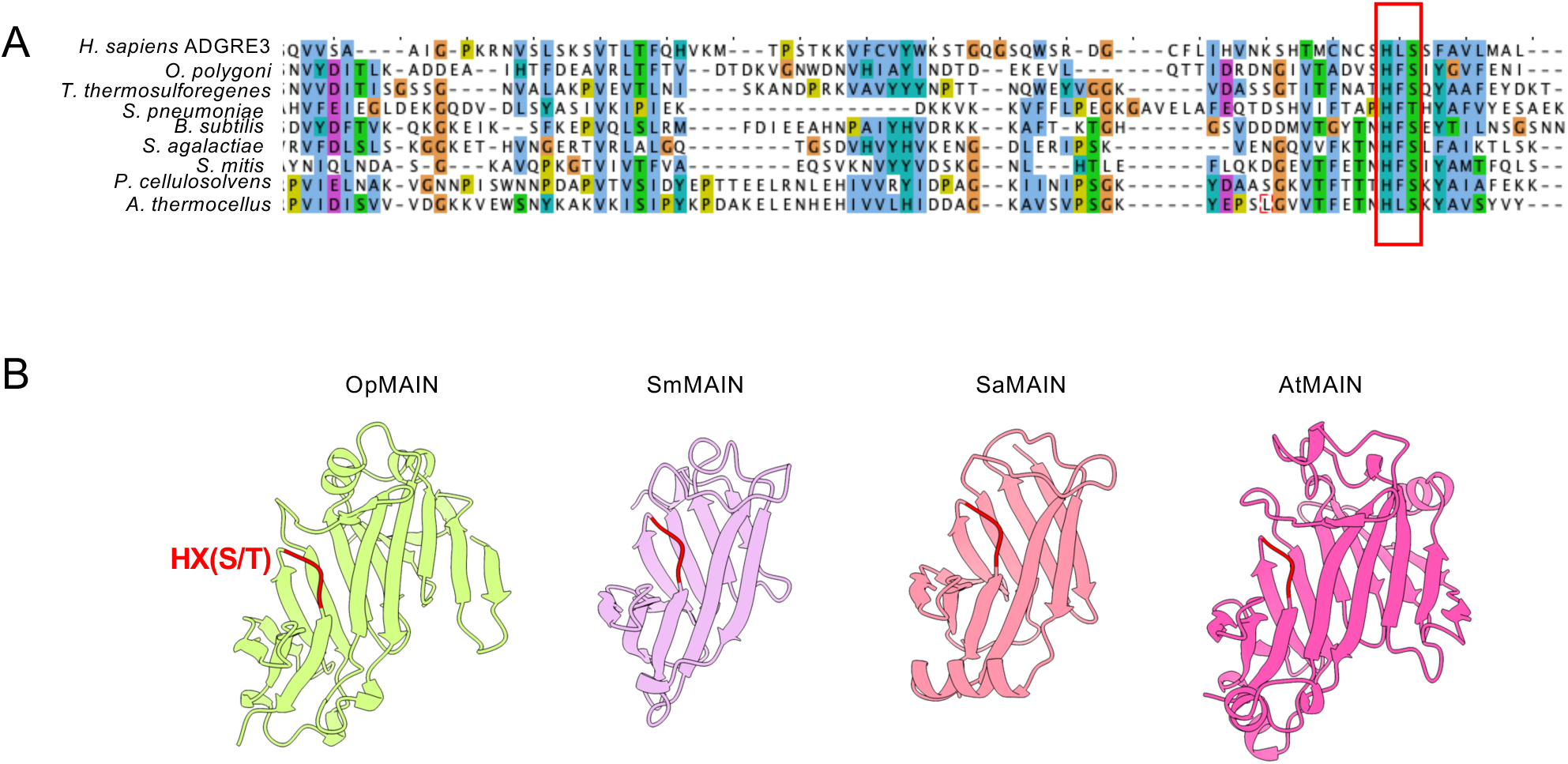
The site of autocleavage is conserved at the sequence and structural level. **(A)** Multiple sequence alignment showing the alignment of the region proximal to the HX(S/T) autoproteolytic site between *H. sapiens* ADGRE3 and several bacterial MAIN domains. **(B)** Alphafold predictions of MAIN domains from the indicated species with the HX(S/T) motif highlighted in red. Confidence metrics for the prediction can be found in Fig. S2.

**Figure S4.**
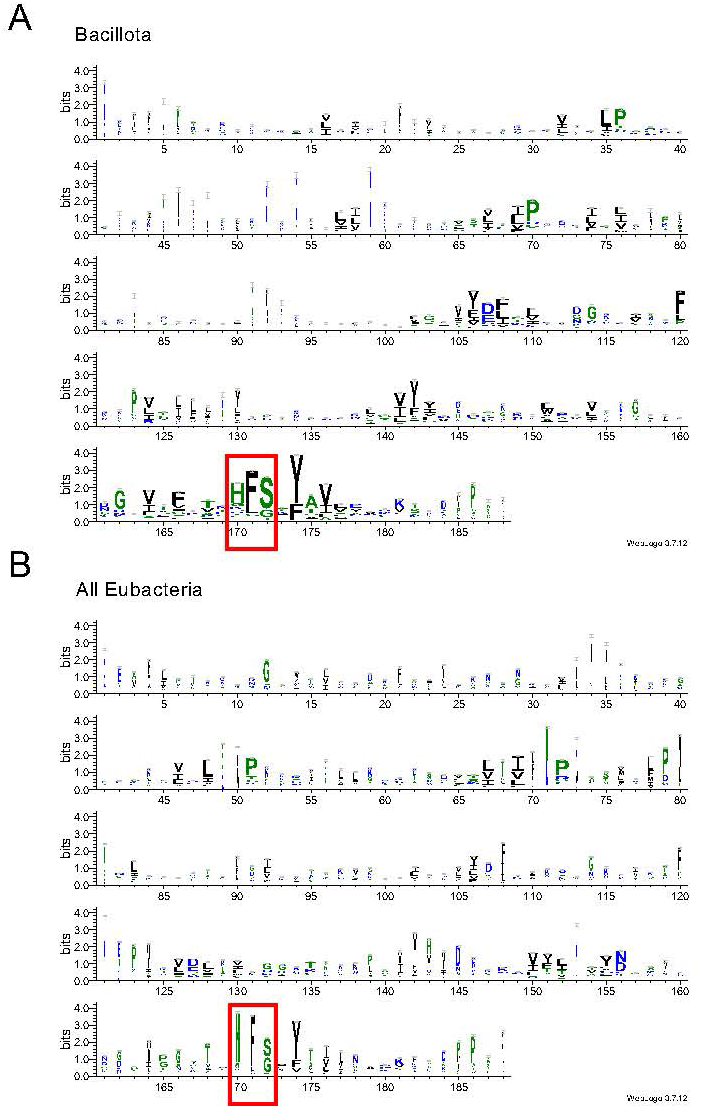
The HFS autocleavage site is largely conserved in Firmicutes. **(A)** Weblogo built from a multiple sequence alignment constructed from MAIN domains found only in Bacillota species. **(B)** Weblogo built from a multiple sequence alignment constructed from MAIN domains found in all Eubacteria and Archaea.

**Figure S5.**
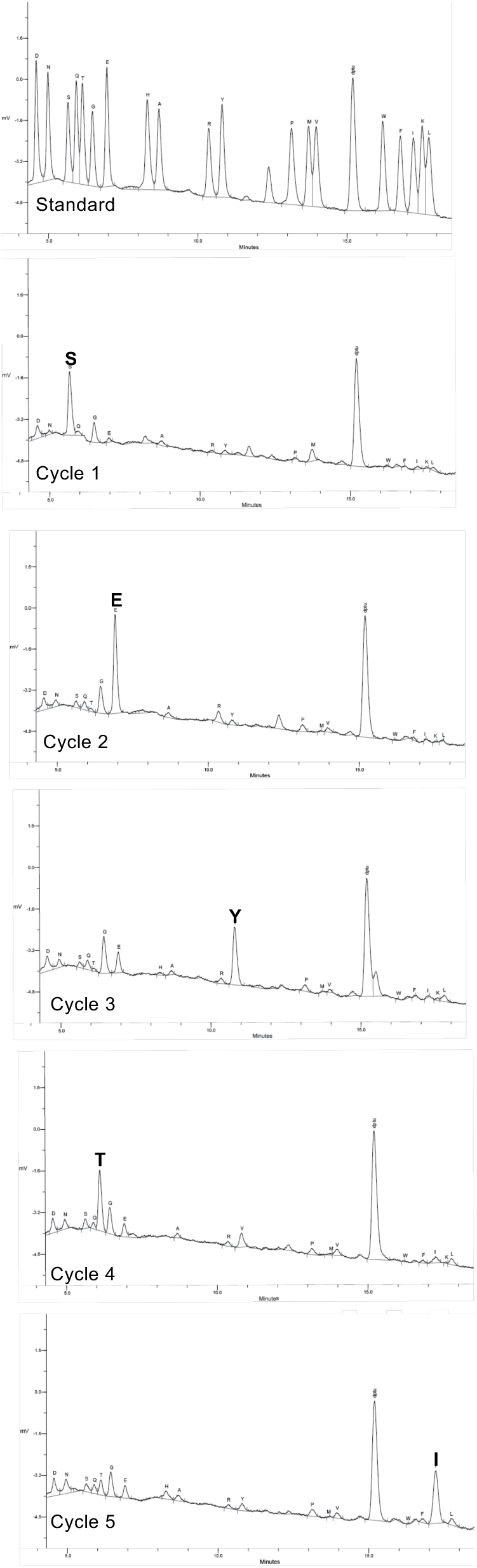
Edman degradation reveals that autocleavage occurs at the conserved HFS motif. Chromatograms from 5 cycles of Edman degradation for the C-terminal cleavage product of BsMAIN as compared to standards of each amino acid.

**Figure S6.**
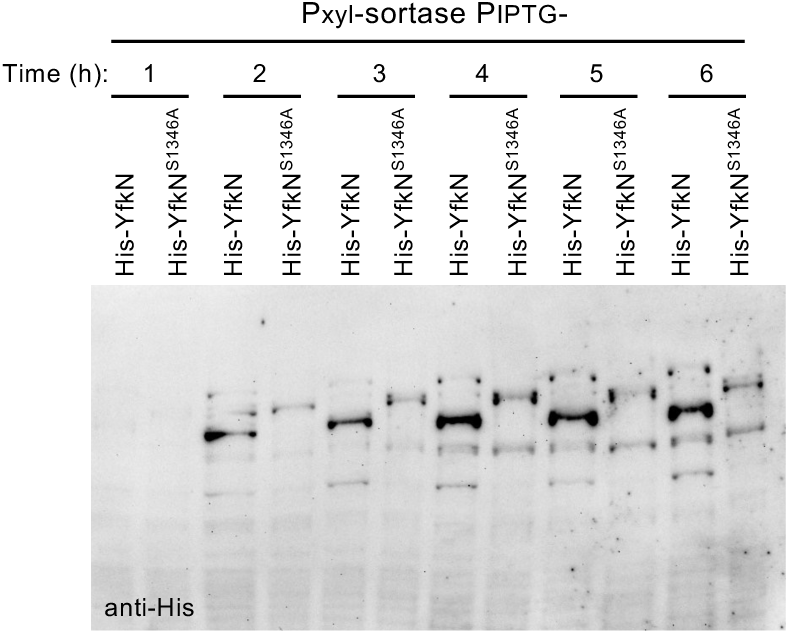
His-YfkN expression increases as cells enter stationary phase. Immunoblot of the indicated *B. subtilis* strains expressing His-YfkN and His-YfkN (S1346A) from a strong IPTG regulated promoter induced with 500 µM IPTG. Cells were subcultured to mid-log, back-diluted to OD600 = 0.05, and grown for 6 hours into stationary phase. Every hour, 1 mL of cells normalized to OD = 1 were harvested, lysed, and analyzed by immunoblot.

**Figure S7.**
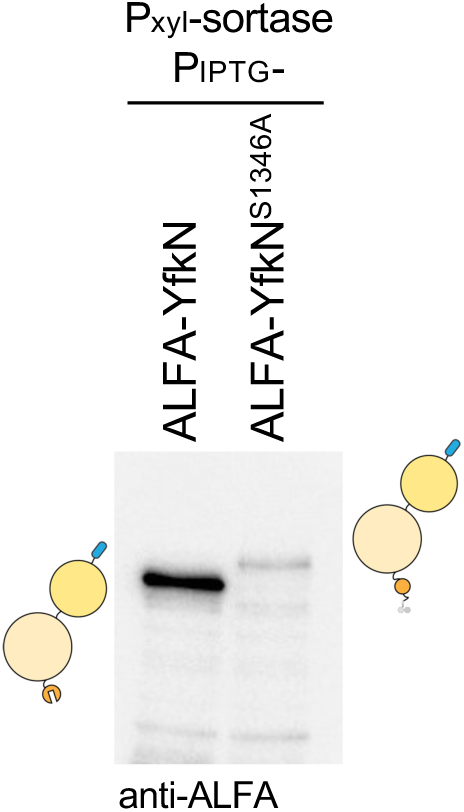
ALFA-tagged YfkN reveals a size shift between wild-type and an autoproteolysis-deficient mutant. Immunoblot of *B. subtilis* stationary phase cells expressing ALFA-YfkN or ALFA-YfkN(S1346A) with 500 µM IPTG. Cells express sortase with 20 mM xylose.

**Figure S8.**
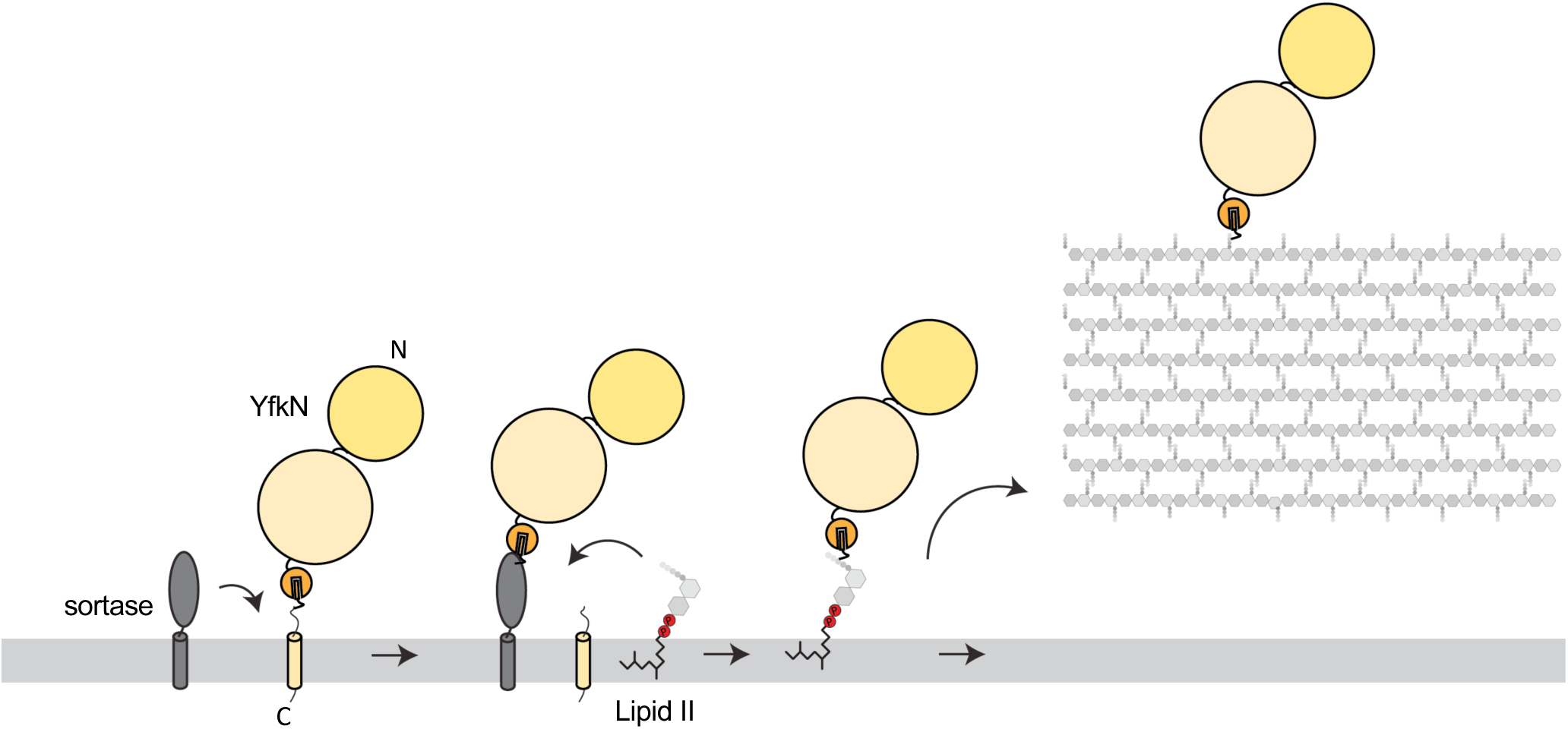
Mechanism of sortase-mediated cell wall anchoring. Sortase substrates have recognition sequences at their C-termini. They are anchored in the membrane by this sequence, prior to transmembrane cleavage and crosslinking to the mDAP residue of the stem peptide of lipid II. Lipid II is then polymerized into the peptidoglycan cell wall.

**Figure S9.**
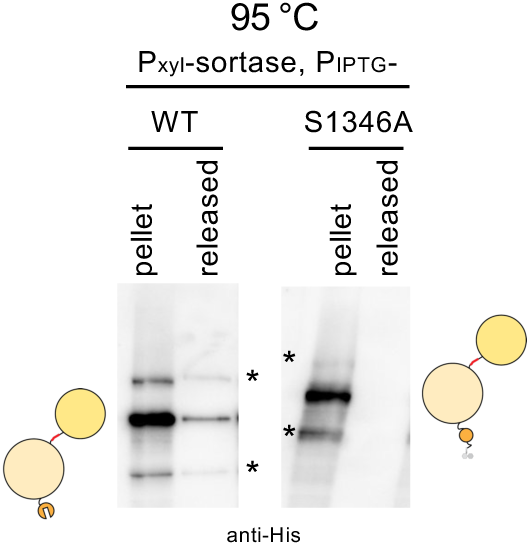
Heat treatment results in YfkN ectodomain release. Immunoblot of *B. subtilis* cells expressing His-YfkN and His-YfkN(S1346A) from a strong IPTG regulated promoter induced with 500 µM IPTG. Cells were subcultured to mid-log, back-diluted to OD600 = 0.05, and grown for 6 hours into stationary phase. Cells were pellet, resuspended in PBS and heated at 95°C for 5 minutes. Cell pellet and supernatant were separated by centrifugation. Anomalous migrating species are indicated (*).

**Figure S10.**
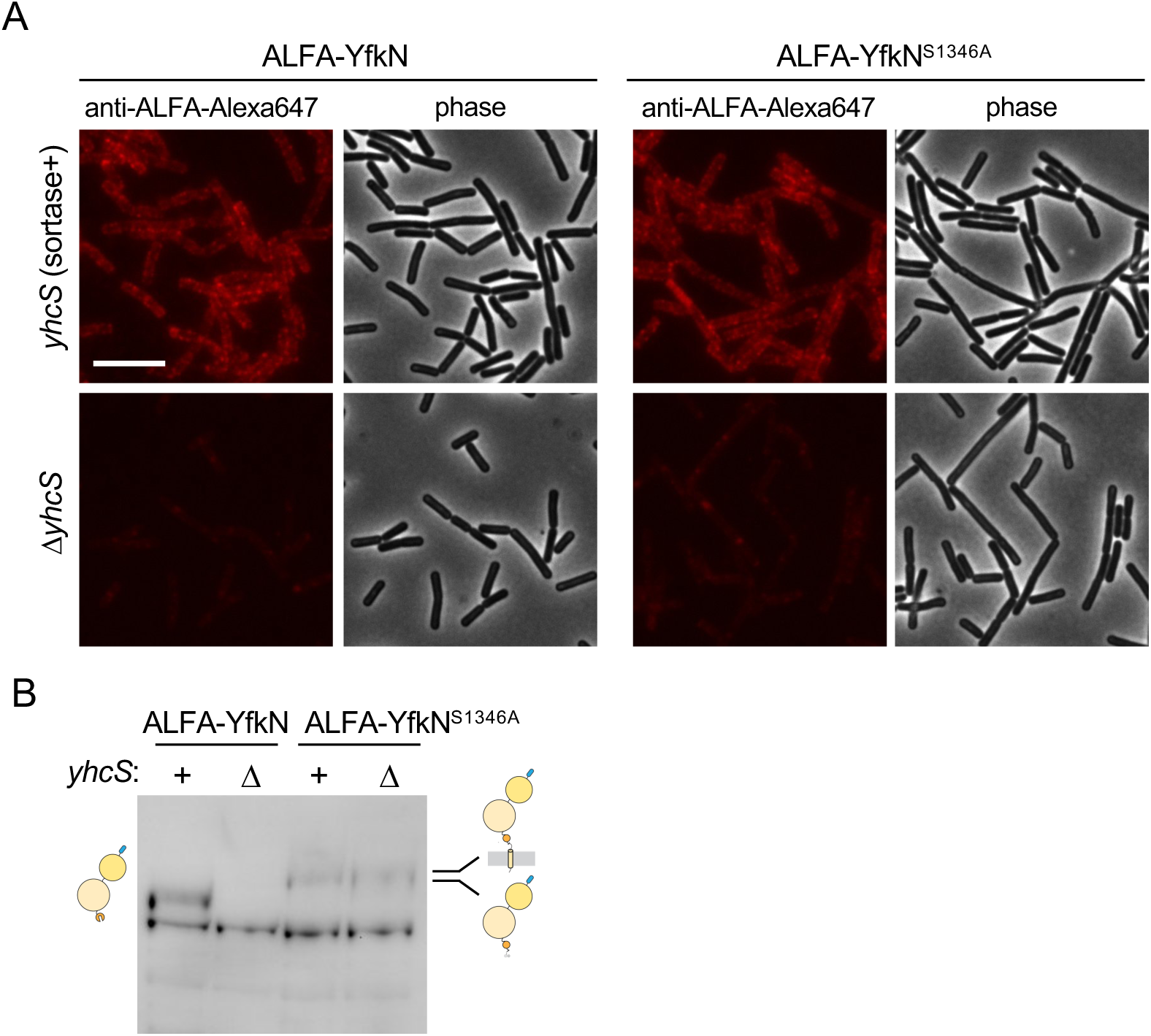
Anti-ALFA-Alexa647 does not label membrane-anchored ALFA-YhcN. **(A)** Fluorescence and phase-contrast micrographs of stationary phase *B. subtilis* cells expressing ALFA-YfkN and ALFA-YfkN(S1346A) in the presence or absence of sortase (YhcS) immunolabeled with anti-ALFA-Alexa647. anti-ALFA-ALexa647 barely labels membrane anchored ALFA-YfkN(S1346A), even though it produced at levels similar to wall-anchored ALFA-YfkN(S1346A), as seen in Figure S12B. YhcN variants were expressed with 500 µM IPTG. Scale bar indicates 5 µm. **(B)** Immunoblot of cells expressing ALFA-YfkN and ALFA-YfkN(S1346A) in the presence (+) or absence (Δ) of sortase (*yhcS*). YfkN is unstable in the absence of sortase, but the autoproteolysis-deficient mutant is stable in the presence and absence of sortase.

**Figure S11.**
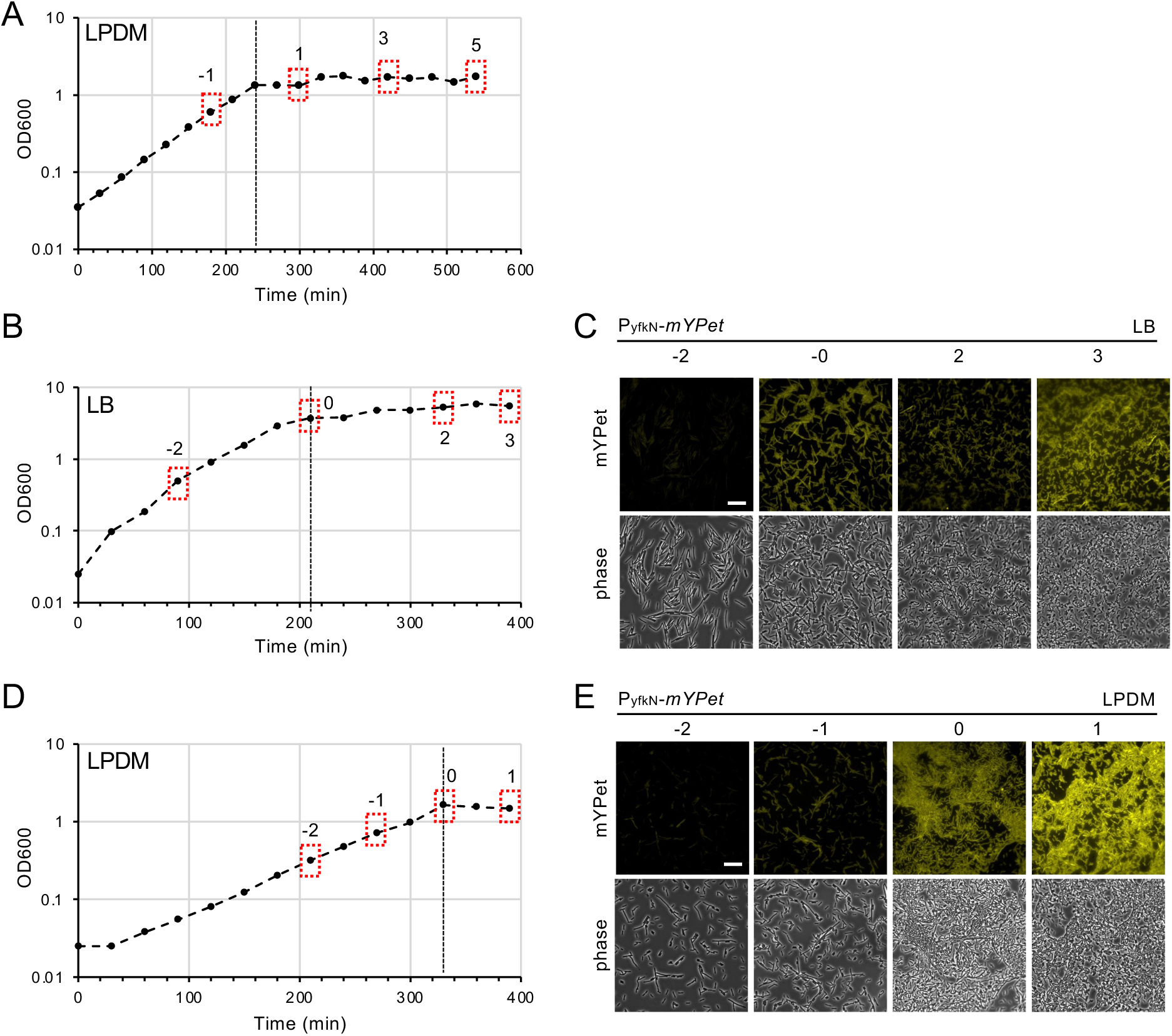
YfkN expression is induced in late stationary phase. **(A)** Growth curve of the time course shown in Figure 3F. **(B)** Growth curve of *B. subtilis* cells harboring the P*_yfkN_*-*mYPet* reporter grown in LB. **(C)** Fluorescence and phase-contrast micrographs of cells at the timepoints indicated in (B). **(D)** Growth curve of *B. subtilis* cells harboring the P_yfkN_-mYpet reporter grown in LPDM. **(E)** Timepoints in (C) & (E) represent time in hours relative to entry into stationary phase. Horizontal dashed lines indicate entry into stationary phase. Scale bar indicates 10 µm.

**Figure S12.**
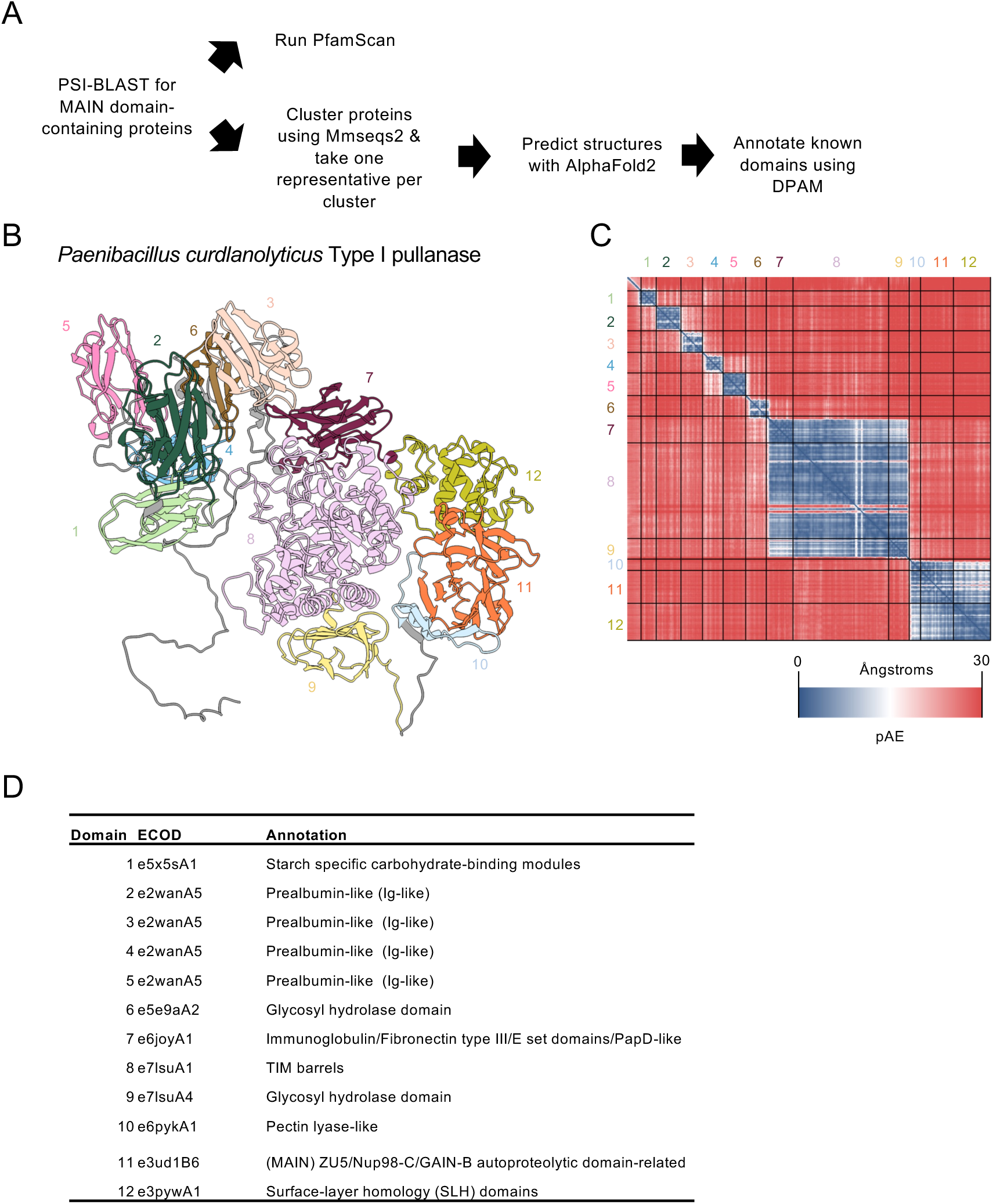
Bioinformatic analysis of MAIN domain-containing proteins. **(A)** 15,000 MAIN domain containing proteins were identified by PSI-BLAST using *B. subtilis* YfkN’s MAIN as a query. The results were used as input for PfamScan, and separately clustered using Mmseqs2 and one representative protein from each cluster used to predict the protein structure by AlphaFold2. Known domains were annotated using DPAM-AI. **(B-D)** DPAM-AI analysis of one representative MAIN domain-containing protein from *Paenibacillus curdlanolyticus* (WP_006036052). (B) Alphafold2-predicted structure of WP_006036052 with domains annotated by DPAM-AI colored. (C) pAE plot of the predicted structure in (B). Black lines indicate domain cutoffs identified by DPAM-AI. (D) Table listing the evolutionary classification of domains (ECOD) domains identified in WP_006036052.

**Figure S13.**
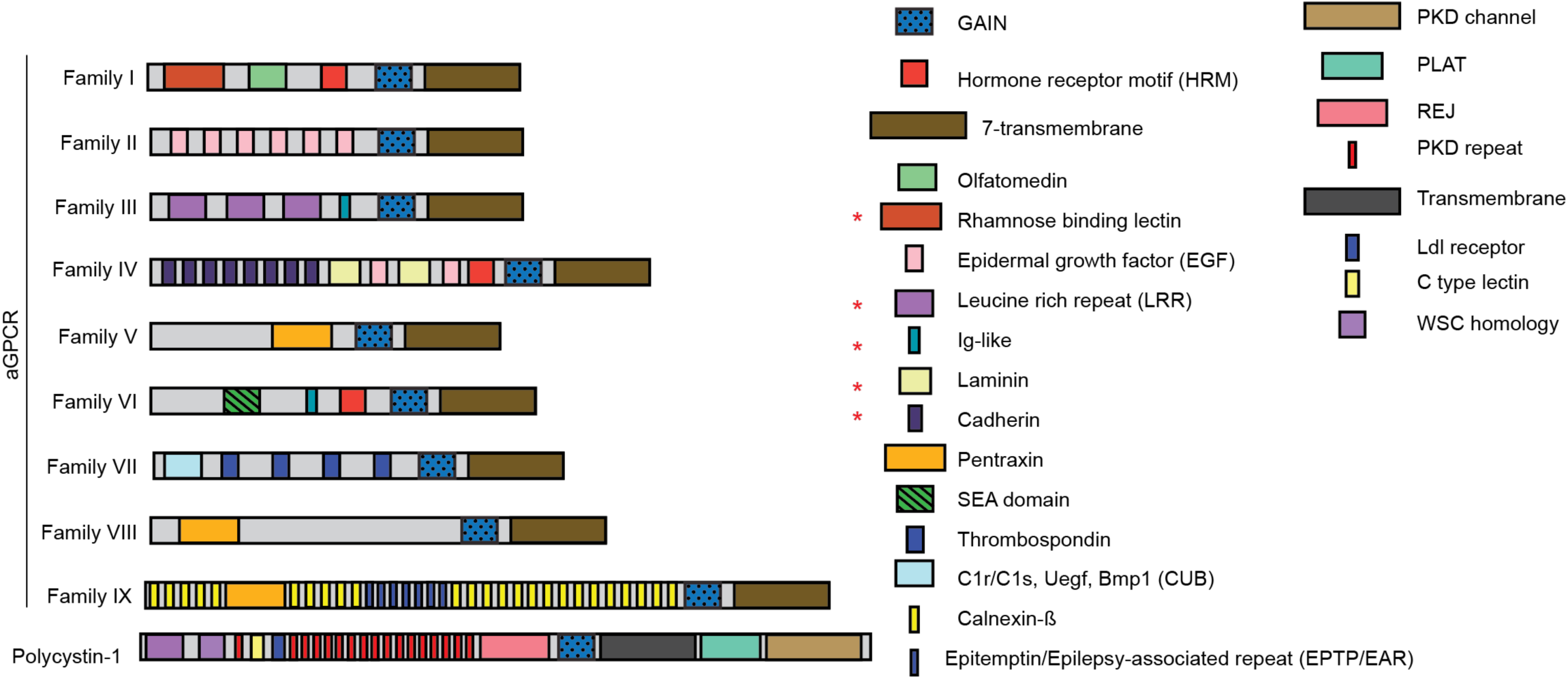
aGPCRs and MAIN-domain containing proteins have common domains. Schematic of aGPCR protein architectures. The 33 aGPCRs can be subdivided into 9 families, which have similar domains. Red asterisks indicate domains that are found on both aGPCRs and MAIN domain containing proteins. Figure adapted from Langenhan et al (2013).

**Figure S14.**
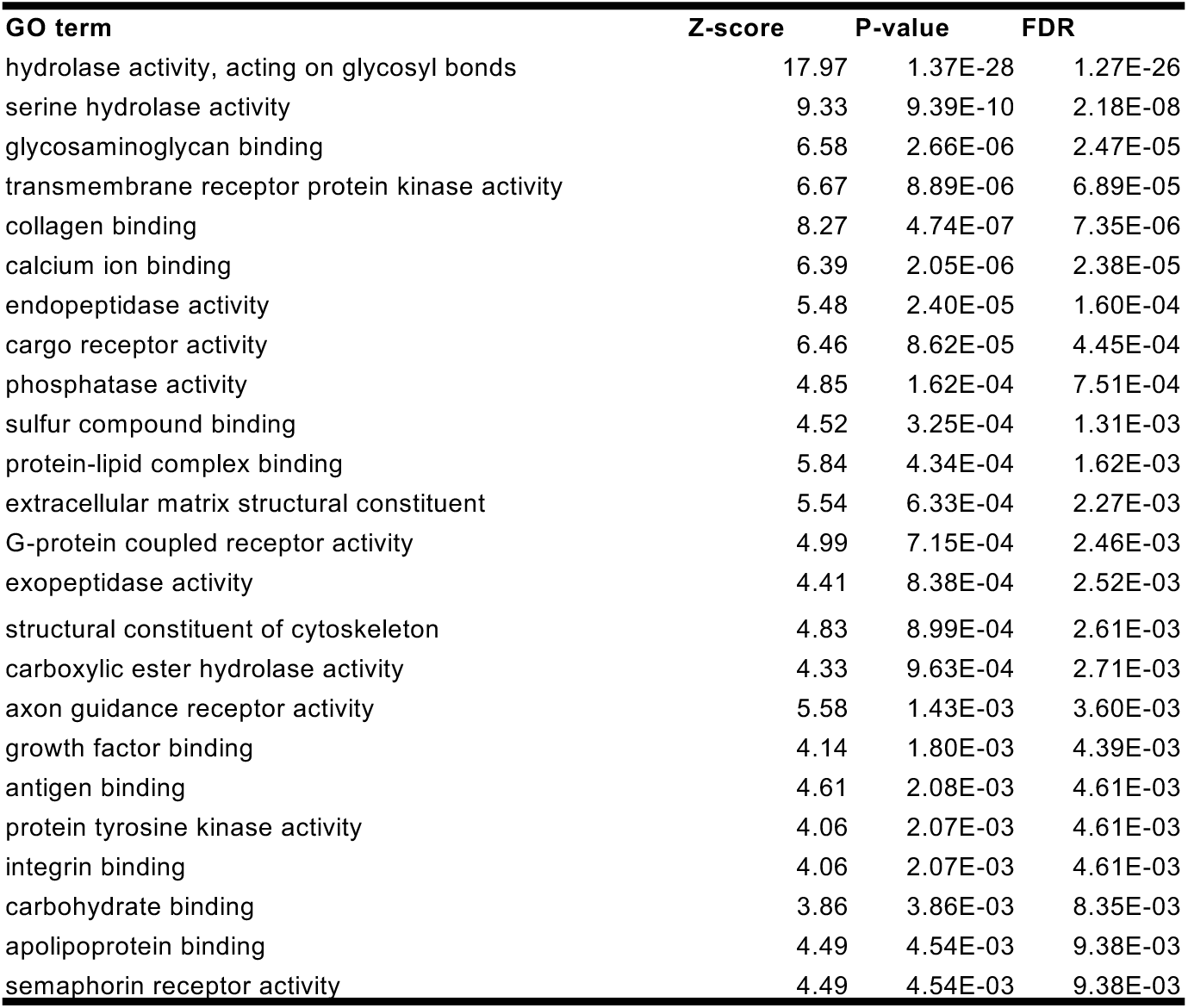
GO enrichment analysis indicates that carbohydrate hydrolase activity is enriched in MAIN-domain containing proteins. A GO enrichment analysis was performed on the Pfam domains encoded on the same polypeptide as MAIN domains. FDR = false discovery rate.

**Figure S15.**
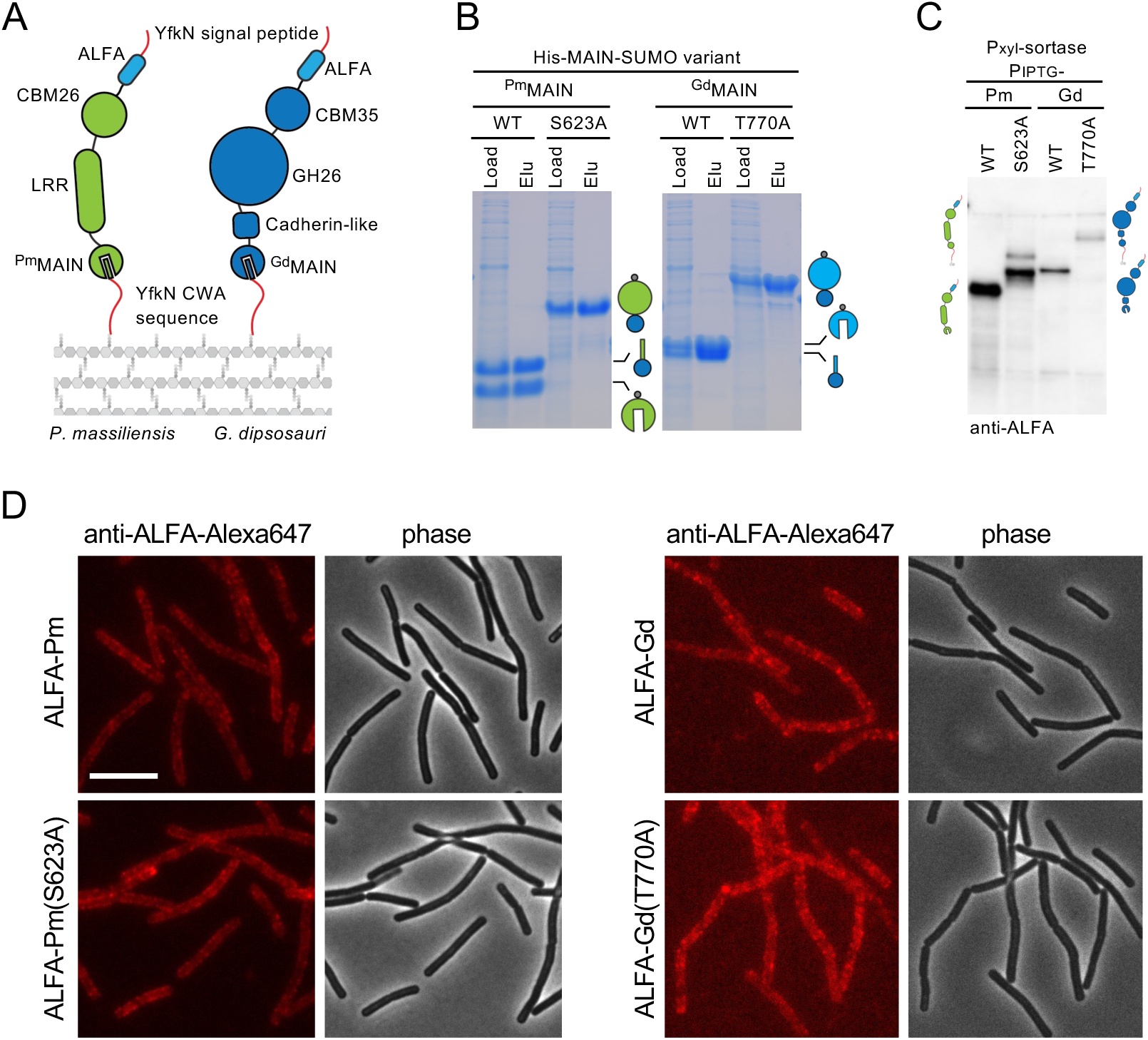
MAIN domain containing proteins from diverse bacteria are autoproteolytic and surface associated. **(A)** Schematic of the system used to anchor chimeric proteins to the *B. subtilis* cell wall. The YfkN signal sequence was fused to the N-terminus and the YfkN cell wall anchoring (CWA) sequence and unstructured region were fused to the C-terminus. MAIN domain containing proteins from *Pseudoruminococcus massiliensis* (WP_106763535) and *Gracilibacillus dipsosauri* (WP_109985763). CBM26 is a putative starch binding fold, CBM35 is putative xylan-binding domain, GH26 is a putative xylan or mannan polysaccharide lyase. Both chimeras were ALFA-tagged at their N-termini just after the signal sequence. **(B)** Coomassie-stained polyacrylamide gel of His-MAIN-SUMO constructs of the indicated variants. Constructs were expressed in *E. coli* and clarified lysate (Load) were incubated with Ni^2+^-NTA resin. Proteins were eluted (Elu) with imidazole. **(C)** Immunoblot of *B. subtilis* cells expressing the indicated constructs and autoproteolysis deficient mutants from (A). **(D)** Fluorescence and phase micrographs of *B. subtilis* cells expressing ALFA-tagged variants of the indicated chimeras. Cells were grown 6 hours into stationary phase and labeled with anti-ALFA-Alexa647. Scale bar indicates 5 µm.

**Figure S16.**
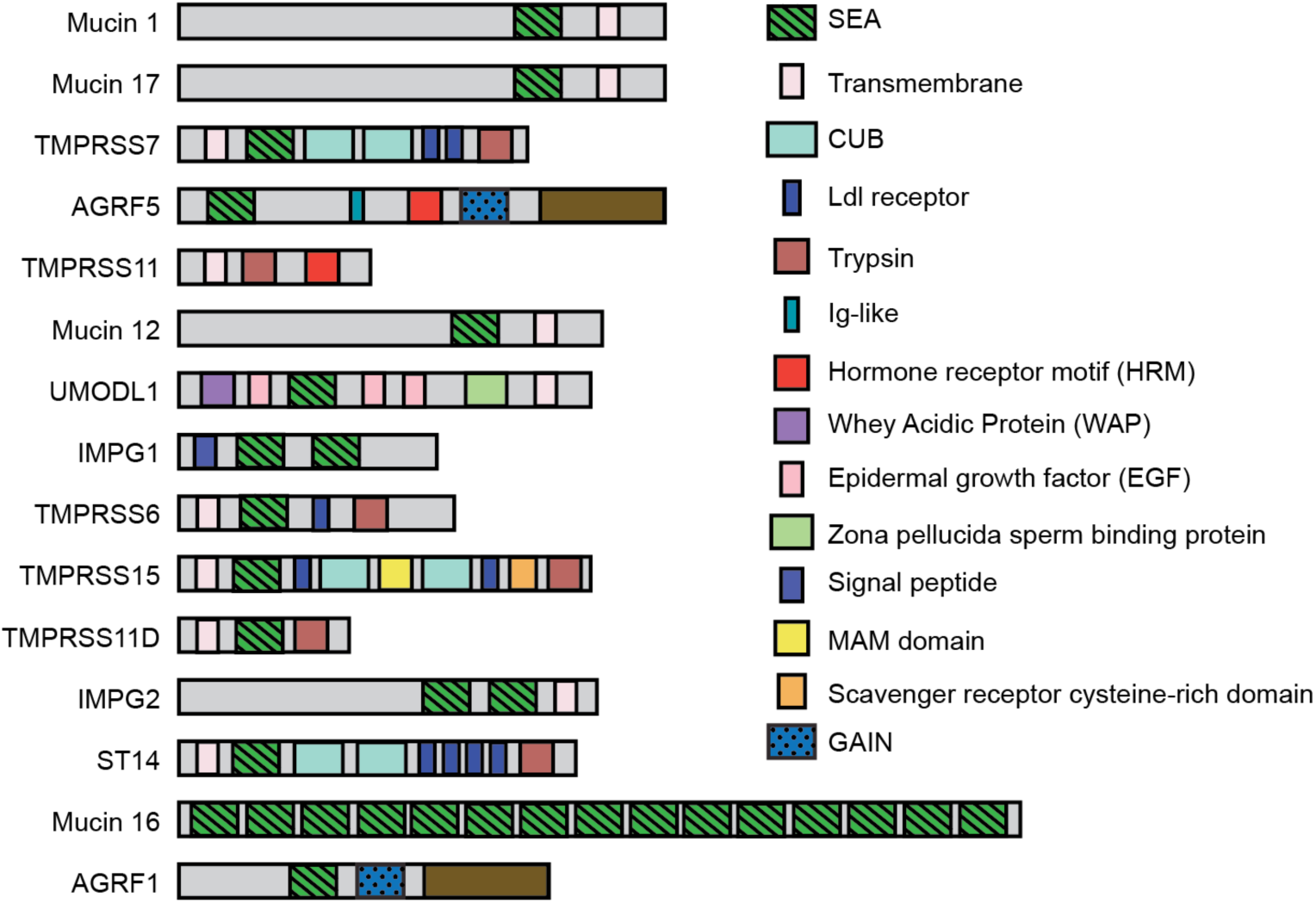
SEA domains are fused to cell surface adhesion domains. Schematic depiction of the SEA domain-containing proteins in the *H. sapiens* proteome. SEA domain-containing proteins were identified by PSI-BLAST and fused domains annotated using PfamScan.

**Figure S17.**
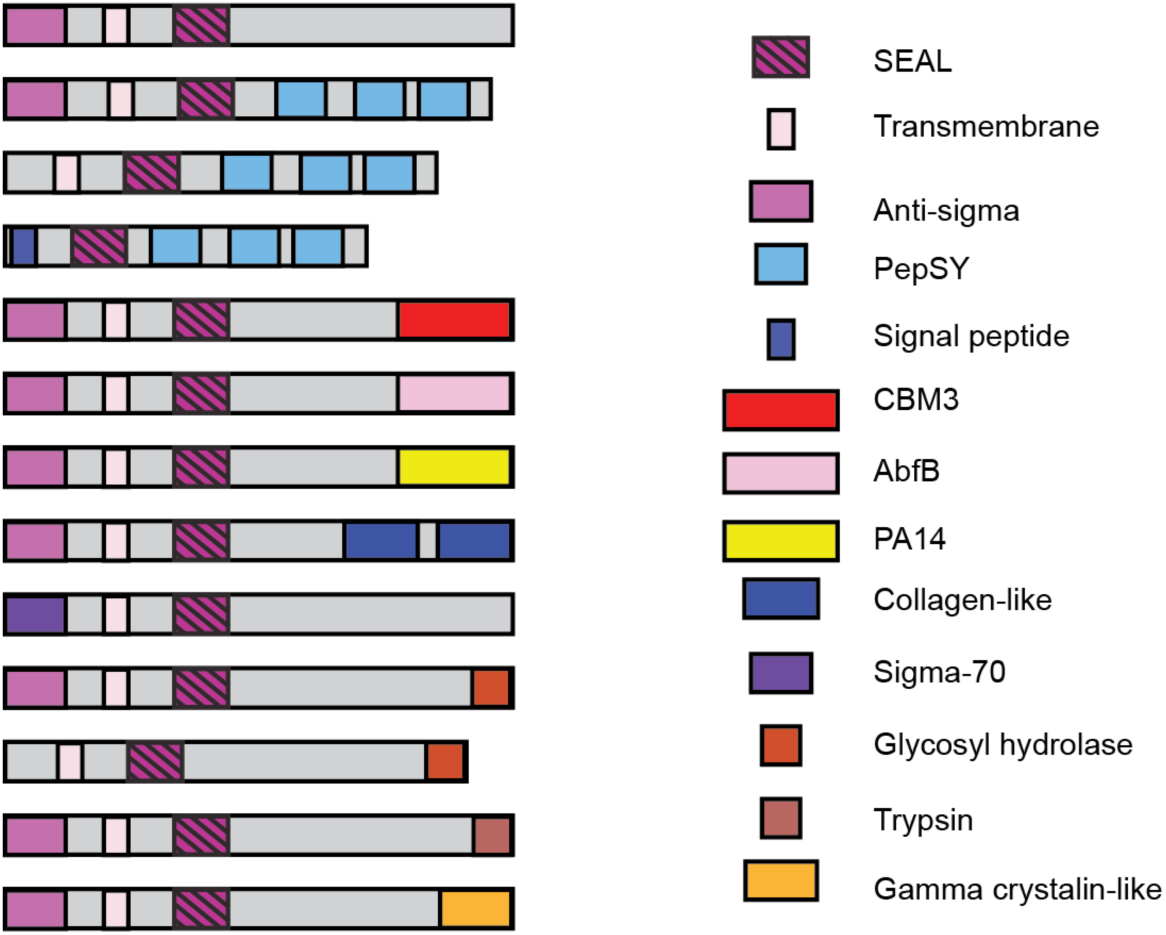
SEA domains are fused to lectins and adhesion domains. Schematic depiction of SEAL domain-containing proteins from diverse bacteria. Ated from Brogan et al.

## Supplemental Table Legends

**Table S1. Pfam domains associated with MAIN domains.** Quantification of Pfam domains found listed in Supplementary Table S2.

**Table S2. Pfam analysis of MAIN domain containing proteins.** Results of the Pfam analysis of ∼15,000 MAIN domain-containing proteins. Domains are listed in columns left to right as predicted from N to C-terminus.

**Table S3. DPAM analysis of MAIN domain containing proteins.** Results of the DPAM analysis of 2,843 representative MAIN domain-containing proteins. Domains are listed in columns left to right as predicted from N to C-terminus. Domains ZU and GPS refer to the GAIN domain fold.

**Table S4. DPAM domains associated with MAIN domains.** Quantification of ECOD domains found listed in Supplementary Table S3.

**Table S5. Strains used in this study. Table S6. Plasmids used in this study.**

**Table S7. Oligonucleotides used in this study. Table S8. GBlocks used in this study.**

**Strain Construction:**

Insertion-deletion mutants in *B. subtilis* were constructed by transforming 3-piece isothermal assemblies containing 1000bp upstream homology, 1000bp downstream homology, and an antibiotic resistance cassette. They were generated using the following oligos: oAB27/28 (universal oligos for antibiotic cassettes), *yfkN* (oAB465/466 & oAB467/468). Ectopic integration was performed by transformation of plasmids linearized by digestion with ScaI.

**Plasmid Construction:**

All plasmids were sequence verified using sanger or nanopore sequencing.

**pAB221 [His-yfkN(MAIN)-SUMO (amp)]**

pAB221 was generated in a 2-piece isothermal assembly reaction using a PCR product amplified from PY79 gDNA using oAB443 & oAB444 and pTD68 [His-SUMO (amp)] digested with NheI.

**pAB222 [His-yfkN(MAIN)-S1346A-SUMO (amp)]**

pAB222 was generated in a KLD (NEB) reaction using a PCR product amplified from pAB221 using oAB436 & oAB437.

**pAB225 [yvbJ-PxylA-yhcS (kan) (amp)]**

pAB225 was generated in a 2-piece isothermal assembly reaction using a PCR product amplified from PY79 gDNA using oAB441 & oAB442 and pCB133 [yvbJ::PxylA (kan) (amp)] digested with XhoI & BamHI.

**pAB227 [yhdG::Phyperspank-yfkN-His6 (spc) (amp)]**

pAB227 was generated in a 3-piece isothermal assembly reaction using PCR products amplified from PY79 gDNA using oAB443 & 448 and oAB444 & 447, and pCB101 [yhdG::Phyperpank (spc) (amp)] digested with HindIII & SpeI.

**pAB228 [yhdG::Phyperspank-yfkN-S1346A-His6 (spc) (amp)]**

pAB228 was generated in a KLD (NEB) reaction using a PCR product amplified from pAB227 using oAB436 & oAB437.

**pAB230 [His-yfkN(MAIN)-S1346T-SUMO (amp)]**

pAB228 was generated in a KLD (NEB) reaction using a PCR product amplified from pAB221 using oAB437 & oAB450.

**pAB233 [His-SpneumoniaeR6_MAIN-SUMO (amp)]**

pAB233 was generated in a 2-piece isothermal assembly reaction using a PCR product amplified from the SpMAIN GBlock using oAB455/456 and pTD68 [His-SUMO (amp)] digested with NheI.

**pAB234 [His-Sagalactiae_MAIN-SUMO (amp)]**

pAB234 was generated in a 2-piece isothermal assembly reaction using a PCR product amplified from the SaMAIN GBlock using oAB455/456 and pTD68 [His-SUMO (amp)] digested with NheI.

**pAB235 [His-Athermocellus_MAIN-SUMO (amp)]**

pAB235 was generated in a 2-piece isothermal assembly reaction using a PCR product amplified from the AtMAIN GBlock using oAB455/456 and pTD68 [His-SUMO (amp)] digested with NheI.

**pAB238 [Opolygoni_MAIN-SUMO (amp)]**

pAB238 was generated in a 2-piece isothermal assembly reaction using a PCR product amplified from the OpMAIN GBlock using oAB455/456 and pTD68 [His-SUMO (amp)] digested with NheI.

**pAB239 [Smitis_MAIN-SUMO (amp)]**

pAB239 was generated in a 2-piece isothermal assembly reaction using a PCR product amplified from the SmMAIN GBlock using oAB455/456 and pTD68 [His-SUMO (amp)] digested with NheI.

**pAB240 [His-Sagalactiae_MAIN-S115A-SUMO (amp)]**

pAB222 was generated in a KLD (NEB) reaction using a PCR product amplified from pAB239 using oAB463 & oAB464.

**pAB247 [yhdG::Phyperspank-ALFA-yfkN (spc) (amp)]**

pAB247 was generated in a 3-piece isothermal assembly reaction using a PCR product amplified from PY79 gDNA using oAB443 & 482, a PCR product amplified from PY79 gDNA using oAB444 & oAB483, and pCB101 [yhdG::Phyperspank (spc)] digested with HindIII & SpeI.

**pAB248 [yhdG::Phyperspank-ALFA-yfkN-S1346A (spc) (amp)]**

pAB247 was generated in a 3-piece isothermal assembly reaction using a PCR product amplified from PY79 gDNA using oAB443 & 482, a PCR product amplified from BAB1144 [yfkN-S1346A] gDNA using oAB444 & oAB483, and pCB101 [yhdG::Phyperspank (spc)] digested with HindIII & SpeI.

**pAB259 [sacA::yfkN-mYPet (phleo) (amp)]**

pAB259 was generated in a 3-piece isothermal assembly reaction using a PCR product amplified from Bs168 gDNA using oAB493 & oAB494, a PCR product amplified from pAB139 [ycgO::Pveg-ydaO-optRBS-mYPet (erm)] using oAB495 & oAB343, and pNC15 [sacA::(phleo)] digested with HindIII & BamHI.

**pAB261 [pminiMAD yfkN-S1346A]**

pAB261 was generated in a 3-piece isothermal assembly reaction using a PCR product amplified from PY79 gDNA using oAB501 & oAB502, a PCR product generated from PY79 gDNA using oAB503 & oAB504, and pminiMAD digested with HindIII & EcoRI.

**pAB263 [yhdG::Phy-spank-SS(yfkN)-His6-NotI-CWA(yfkN) (spc) (amp)]**

pAB263 was generated using a 3-piece isothermal assembly reaction using a PCR product amplified from PY79 gDNA using oAB443 & oAB512, a PCR product amplified from PY79 gDNA using oAB513 & oAB444, and pCB101 [yhdG::Phyperspank (spc) (amp)] digested with HindIII & SpeI.

**pAB264 [yhdG::Phy-spank-SS(yfkN)-PmCBM26-CWA(yfkN) (spc) (amp)]**

pAB264 was generated in a 2-piece isothermal assembly reaction using a PCR product amplified from the PmCBM26 GBlock using oAB514 & oAB515, and pAB263 digested with NotI.

**pAB265 [yhdG::Phy-spank-SS(yfkN)-PmCBM26-CWA(yfkN)-S623A (spc) (amp)]**

pAB265 was generated in a 3-piece isothermal assembly reaction using a PCR product amplified from the PmCBM26 GBlock using oAB514 & oAB552, a PCR product amplified from pAB264 using oAB551 & oAB444, and pAB263 digested with NotI.

**pAB269 [His-PmCBM26(MAIN)-SUMO (amp)]**

pAB269 was generated in a 2-piece isothermal assembly reaction using a PCR product amplified from the PmCBM26 GBlock using oAB525 & oAB526, and pTD68 [His-SUMO (amp)] digested with NheI.

**pAB270 [His-PmCBM26(MAIN)-S623A-SUMO (amp)]**

pAB269 was generated in a 2-piece isothermal assembly reaction using a PCR product amplified from the PmCBM26 GBlock using oAB525 & oAB543, and pTD68 [His-SUMO (amp)] digested with NheI.

**pAB285 [yhdG::Phy-spank-SS(yfkN)-ALFA-PmCBM26-CWA(yfkN) (spc) (amp)]**

pAB285 was generated in a 2-piece ITA using a PCR product amplified from PY79 gDNA using oAB443 & oAB482 and a PCR product amplified from pAB264a [yhdG-Phy-spank-SS(yfkN)-PmCBM26-CWA(yfkN) (spec) (amp)] using oAB562 & oAB444.

**pAB286 [yhdG::Phy-spank-SS(yfkN)-ALFA-PmCBM26-CWA(yfkN)-S623A (spc) (amp)]**

pAB286 was generated in a 2-piece ITA using a PCR product amplified from PY79 gDNA using oAB443 & oAB482 and a PCR product amplified from pAB265a [yhdG-Phy-spank-SS(yfkN)-PmCBM26-CWA(yfkN)-S623A (spec) (amp)] using oAB562 & oAB444.

**pAB288 [yhdG::Phy-spank-SS(yfkN)-ALFA-NotI-CWA(yfkN) (spc) (amp)]**

pAB288 was generated in a 3-piece isothermal assembly reaction using a PCR product amplified from PY79 gDNA using oAB443 & oAB482, a PCR product amplified from PY79 gDNA using oAB564 & oAB444, and pCB101 [yhdG::Ph-spank (spc) (amp)] digested with HindIII & SpeI.

**pAB289 [yhdG::Phy-spank-SS(yfkN)-ALFA-CBM35-CWA(yfkN) (spc) (amp)]**

pAB289 was generated in a 2-piece ITA using a PCR product amplified from the CBM35 GBlock (WP_109985763) using oAB565 & oAB566 and pAB288 [yhdG::Phy-spank-SS(yfkN)-ALFA-NotI-CWA(yfkN) (spc) (amp)] digested with NotI.

**pAB290 [yhdG::Phy-spank-SS(yfkN)-ALFA-CBM35-CWA(yfkN)-T770A (spc) (amp)]**

pAB290 was generated in a Quikchange reaction (Agilent) using a PCR product amplified with Pfu turbo using oAB577 & oAB578.

**pAB295 [His-CBM35(MAIN)-SUMO (amp)]**

pAB295 was generated in a 2-piece isothermal assembly reaction using a PCR product amplified from the CBM35 GBlock (WP_109985763) using oAB571 & oAB572, and pTD68 [His-SUMO (amp)] digested with NheI.

**pAB296 [His-CBM35(MAIN)-T770A-SUMO (amp)]**

pAB296 was generated in a 3-piece isothermal assembly reaction using a PCR product amplified from the CBM35 Gblock (WP_109985763) using oAB571 & oAB578, a PCR product amplified from the CBM35 Gblock (WP_109985763) using oAB572 & oAB577, and pTD68 [His-SUMO (amp)] digested with NheI.

